# tRNA modification profiling reveals epitranscriptome regulatory networks in *Pseudomonas aeruginosa*

**DOI:** 10.1101/2024.07.01.601603

**Authors:** Jingjing Sun, Junzhou Wu, Yifeng Yuan, Leon Fan, Wei Lin Patrina Chua, Yan Han Sharon Ling, Seetharamsing Balamkundu, Dwijapriya, Hazel Chay Suen Suen, Valérie de Crécy-Lagard, Agnieszka Dziergowska, Peter C. Dedon

**Affiliations:** Department of Biological Engineering, Massachusetts Institute of Technology, Cambridge, MA 02139 USA; Antimicrobial Resistance Interdisciplinary Research Group, Singapore-MIT Alliance for Research and Technology, 138602 Singapore; Department of Microbiology and Cell Science, University of Florida, Gainesville, FL 32611 USA; School of Biological Sciences, Nanyang Technological University, Singapore; Department of Food, Chemical & Biotechnology, Singapore of Institute of Technology, 138683 Singapore; Genetic Institute, University of Florida, Gainesville, FL 32611 USA; Institute of Organic Chemistry, Lodz University of Technology, 90-924 Poland

## Abstract

Transfer RNA (tRNA) modifications have emerged as critical posttranscriptional regulators of gene expression affecting diverse biological and disease processes. While there is extensive knowledge about the enzymes installing the dozens of post-transcriptional tRNA modifications – the tRNA epitranscriptome – very little is known about how metabolic, signaling, and other networks integrate to regulate tRNA modification levels. Here we took a comprehensive first step at understanding epitranscriptome regulatory networks by developing a high-throughput tRNA isolation and mass spectrometry-based modification profiling platform and applying it to a *Pseudomonas aeruginosa* transposon insertion mutant library comprising 5,746 strains. Analysis of >200,000 tRNA modification data points validated the annotations of predicted tRNA modification genes, uncovered novel tRNA-modifying enzymes, and revealed tRNA modification regulatory networks in *P. aeruginosa*. Platform adaptation for RNA-seq library preparation would complement epitranscriptome studies, while application to human cell and mouse tissue demonstrates its utility for biomarker and drug discovery and development.

## Introduction

The more than 170 post-transcriptional RNA modifications comprising the epitranscriptome play a crucial role in regulating mRNA translation in all forms of life. Defects in RNA-modifying enzymes drive dozens of diverse human diseases such as cancer, neurological disorders, and metabolic diseases^1–3^, while RNA modifications also play a role in microbial pathogenesis and antimicrobial resistance^4–7^. There is a growing appreciation for the complexity of mechanisms linking transfer RNA (tRNA) modifications to normal and pathological cell phenotypes, such as tRNA reprogramming and codon-biased translation in cell stress response and disease^5, 6, 8–10^. These transcendent behaviors require multi-omic tools to define molecular connections between upstream environmental sensing and signaling pathways that regulate tRNA-modifying enzymes and the tRNA pool and downstream phenotypic changes in cell physiology and pathology. This kind of systems-level information is critical for validating the dozens of RNA-modifying enzymes as a novel class of drug targets^11–13^.

The major hindrance to systems-level analyses of the tRNA epitranscriptome lies in the lack of technology for high-throughput (HT) tRNA modification and tRNA pool analysis^14^. Such technology is needed, for example, to screen for tRNA-related translational defects in the 2000 DepMap cell lines comprising dozens of different cancers^15^, to screen thousands of strains in gene knockout libraries, such as those for *Escherichia coli*^16^, *Bacillus subtilis*^17^, *Pseudomonas aeruginosa*^18^, and *Enterococcus faecalis*^19^, or for whole-cell phenotypic screening of drug libraries for effects on tRNA modification levels. The technology limitations hindering such studies start with tRNA isolation from cells and tissues. Traditional RNA purification methods relying on phenol-based liquid phase extraction are not only difficult to adapt to automated platforms but also fail to sufficiently resolve small RNAs, such as tRNA and miRNA, from total RNA. This necessitates additional size-based separation methods for further isolation of tRNA^20^. The alternative of silica-based spin column methods to isolate small RNA directly from cell lysates is cost-prohibitive, labor-intensive, and time-consuming for large-scale studies.

Even with purified tRNA in hand, it is similarly challenging to quantify modified ribonucleosides in tRNA hydrolysates by automated methods. Here there is a clear advantage to using chromatography-coupled tandem quadrupole mass spectrometry (LC-MS/MS), with recent advances in chromatography reducing run times from 20-30 minutes per sample^20^ down to 9 minutes^21^, for example. A major hindrance, though, lies in the pipeline for signal processing and data mining, which requires significant customization.

Here we report a robust, HT tRNA modification analysis platform involving magnetic bead-based purification of tRNA directly from cell and tissue lysates, rapid LC-MS/MS analysis of ribonucleosides, and a data processing and analysis pipeline. The small RNA purification leg of the platform was validated on broad range of biological samples, including bacteria (*P. aeruginosa*), mammalian cells (Hela, HEK293T) and animal tissue (mouse brain). The combined RNA purification and LC-MS/MS modification features of the platform were then used to screen a *P. aeruginosa* UCBPP-PA14 transposon insertion mutant library consisting of 5,746 mutant strains covering 4,360 non-essential genes^18^. The loss of two dozen known tRNA modification genes and their expected modification products further validated the platform and the results allowed annotation of tRNA modification genes in PA14. More importantly, the screen revealed hundreds of genes affecting tRNA modification levels, with these genes forming regulatory networks for iron-sulfur cluster synthesis and repair, redox homeostasis, amino acid synthesis, and tRNA modifying enzyme co-factor synthesis, among others. The results provide a comprehensive view of the regulatory landscape of tRNA modifications in *P. aeruginosa* and demonstrate the utility of the HT tRNA analytical platform.

## Results

### Developing the high-throughput platform for tRNA modification profiling

The workflow for the HT tRNA analytical platform is shown in **Figure 1A** and begins with growing cells (∼1.6 x 10^8^ CFU PA14, 5 x 10^5^ HeLa cells) or placing tissue samples (10 mg) in wells of a 96-well plate. The samples are then subjected to cell lysis, removal of large RNA and genomic DNA, purification of small RNA, hydrolysis of RNA to ribonucleosides, LC-MS/MS analysis, and signal processing (**Figure 1A**), with each step optimized as detailed in **Supplementary Information**. Bacterial and human cells were all lysed using a buffer containing 4 M guanidine isothiocyanate (GITC) (**Supplementary Figure 1**) with shaking but without enzymatic or mechanical disruption. Animal tissues were lysed with the same buffer supplemented with mechanical disruption using a tissue lyser (**Supplementary Figure 2**). This method facilitated effective inhibition of RNases and other RNA-modifying enzyme activity and further enhanced the size selection resolution of magnetic beads for small RNA separation (**Supplementary Figure 3B**).

**Figure 1.**
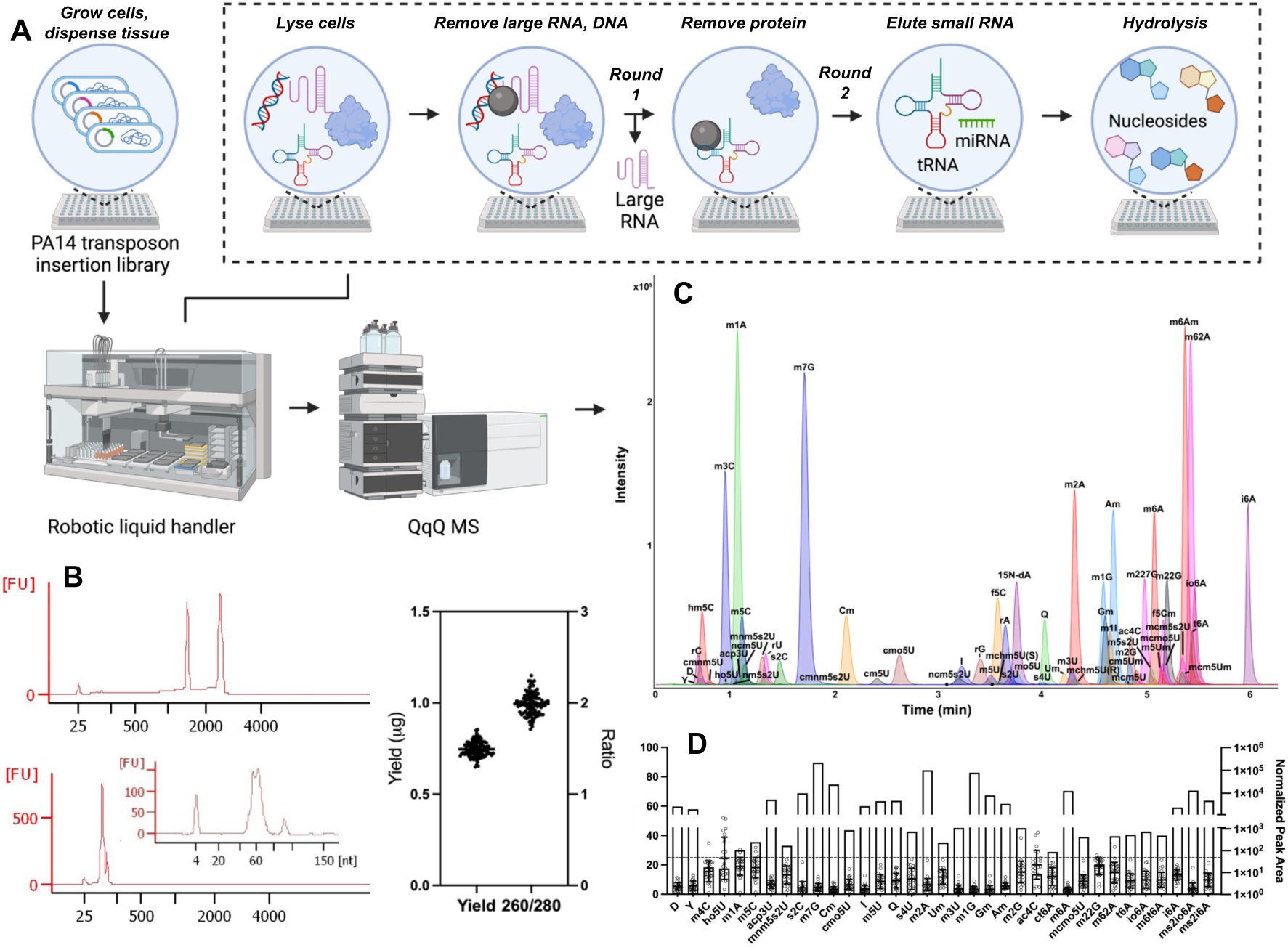
Workflow and validation of tRNA modification profiling platform applied to a PA14 mutant library. (**A**) Cell growth, lysis, and small RNA purification in 96-well plate format, with a two-step magnetic beads-based size selection strategy for tRNA purification direct from crude lysate. All steps are performed with a robotic liquid handler. (**B**) Characterization of purified RNA fractions from PA14 crude lysate (0.3 OD_600_ cells/well). The Bioanalyzer tracings show the quality of purified large RNAs (upper) and small RNAs (lower) assessed using a “pico chip”. Inset: a “small RNA chip” further detailing the small RNA quality. The graph shows the small RNA yield of 747±42 ng/well with an A_260_/A_280_ purity ratio of 1.99±0.12 for n=96. (**C**) An extracted ion chromatogram shows the fast UHPLC-MS/MS method for quantifying 63 modified ribonucleosides (based on synthetic standards) using the dynamic multiple reaction mode (DMRM) of QQQ. (**D**) Quantitative performance of the LC-MS/MS method was evaluated using data from 24 PA14 mutants. The inter-day coefficient of variation (CV) for quantification of 35 RNA modifications was calculated from the average peak area for each RNA modification normalized by the total UV absorbance of the 4 canonical ribonucleosides (see Methods). The dashed line indicates a CV of 25%.

We next used a two-step magnetic beads-based method to isolate small RNA species from crude lysates in 96-well plate format (**Figure 1A**). In the first step (Round 1), RNA binding buffer I containing magnetic beads, salts, and molecular crowding agent (PEG 8000) was added to capture genomic DNA and large RNAs (>150 nt, mainly rRNA and mRNA), leaving small RNAs (<150 nt, mainly tRNA) in the supernatant. In the second step (Round 2), RNA binding buffer II containing isopropanol and fresh beads was added to the supernatant to capture small RNAs. Beads from both rounds 1 and 2 were washed and the nucleic acids eluted as described in **Methods**. This protocol was optimized for the composition of RNA binding buffers I and II (salts, pH, crowding reagents; **Supplementary Figure 3**) to maximize the yield and purity of the small RNAs. In our experiments, both silica- and carboxyl-coated magnetic beads were effective for bacterial and mammalian cells, but we settled on carboxyl-coated beads due to their compatibility with tissues (**Supplementary Figure 3**). The method proved to be as effective as silica column-based commercial kits in size resolution and tRNA yield (**Supplementary Figure 3C**).

The cell lysis and tRNA purification steps were then adapted to a 96-well plate format and automated using a robotic liquid handler (Tecan EVO150). The Tecan workflow includes 9 steps for the tRNA purification process in 1 h for 96 samples (**Supplementary Figure 4**). To ensure consistent and robust results (**Figure 1B, Supplementary Figure 5**), we empirically determined the optimal labware and parameters for aspiration, dispensing, and mixing (**Supplementary Table 1**). For instance, the “Wash buffer residue” step was optimized and conducted twice to thoroughly eliminate ethanol residue, which addressed the issue of peak shape broadening during LC-MS/MS analysis (**Supplementary Figure 6A**). Another important issue was removal of rRNA contamination as carryover from the 1^st^ round of beads. We introduced a plate centrifugation action between the two magnet pull-downs to ensure efficient removal of 1^st^ round beads (**Supplementary Figure 6B**). Using the automated approach, the average yield of small RNA from approximately 0.3 OD_600_ *Pseudomonas* cells is 747±42 ng (n=96) with an average 260/280 nm ratio of 1.99±0.12 (**Figure 1B**). Collectively, this method provided high-quality tRNA samples at low cost ($0.3 per sample) and high efficiency (1 h for 96 samples).

The second portion of the platform, LC-MS/MS analysis of RNA modifications, required optimization of both the HPLC resolution and the MS/MS quantification of ribonucleosides to increase sensitivity and reduce the typical run time of 20-30 minutes for one sample.^20^ Here we coupled a rapid UHPLC method with dynamic multiple reaction monitoring for analysis of >60 RNA modifications in a 6-minute HPLC run (**Figure 1A**). This approach was found to be applicable for analyses of tRNA from bacteria, mammalian cells, and animal tissues (**Supplementary Figure 2**). Of the two compact columns assessed — BEH C18 and HSS T3 —the former outperformed the latter with a higher plate count and a shorter run time (**Supplementary Figure 7J**). Both the LC configurations (**Supplementary Figure 7**) and MS parameters (**Supplementary Table 2**) were fine-tuned to optimize identification and quantification of modifications, including isobaric methylation isomers (**Supplementary Figure 7**). The sensitivity of the method was confirmed with limits of detection and quantification for modified ribonucleosides in the low femtomole range, including problematic U modifications (**Supplementary Table 3**). Method performance was further assessed using tRNA from wild-type PA14, focusing on retention time stability, chromatographic peak characteristics, and signal carryover. The standard deviation for retention times was less than 5 seconds (**Supplementary Figure 8A**), with coefficient of variation for peak area less than 10% (**Supplementary Figure 8B**). The full width at half maximum (FWHM) for most ribonucleosides stayed under 5 seconds, only cmo^5^U extended to 7 seconds (**Supplementary Figure 8C**). No negligible sample carryover was observed throughout the spectrum (**Supplementary Figure S8D**). To validate the reliability of the rapid LC-MS/MS method, we analysed the same sample matrix (n=16) using both fast (6-minute) and conventional methods and found them to be strongly correlated (**Supplementary Figure 8F**; Pearson correlation r = 0.927, *p* < 0.0001). In addition, the composition of enzyme cocktails for tRNA hydrolysis was examined based on previous work in our lab (**Supplementary Figure 9A**)^22^. An adenosine deaminase inhibitor, here coformycin, was found to be essential to obtain accurate inosine levels, with deaminase contamination of enzyme preparations evident from increased inosine in the absence of coformycin. However, the cytidine deaminase inhibitor appears unnecessary as no significant m^3^C/m^5^C to m^3^U/m^5^U deamination was detected. Moreover, the inclusion of the iron chelator deferoxamine was proved to protect ho^5^U from iron-induced Fenton reaction^22^. Lastly, we replaced the polyethersulfone 10K spin-filter used in previous protocols with a 0.2 µM stainless steel inline filter to prevent significant loss of hydrophobic ribonucleosides (**Supplementary Figure 9C**).

Finally, the third leg of the platform involves data processing pipeline to manage conversion of hardware-specific signals to normalized signal intensities comparable across different analytical runs, to collate signal intensities with gene names, and to calculate fold-change values relative to the adjusted mean of samples run in 2 h. This mean is calculated based on the expectation that the majority of mutants exhibit insignificant modification level changes. This is substantiated by our screening data, which indicates that over 94% of the 17,2860 measurements display less than 1.2-fold changes. This approach waives the comparison with wild-type strain which is suboptimal served as a control as it cannot be cultured in the same condition/plate as mutants. Additionally, the potential for signal drift is significant^23^, given that LC-MS analysis of a single 96-well plate may require a full day. To mitigate these issues, we use an approach to calculate adjusted mean for fold change calculation. We initially calculated modification levels by averaging UV-normalized peak areas for each row. Then, we eliminated any data points exhibiting more than a two-fold change before recalculating the final average, which then served as the baseline for fold change calculations for each respective row. The details of the data processing are described in **Methods**.

The fully optimized platform allows processing of one 96-well plate per hour for tRNA purification and hydrolysis, followed by one plate processed every 15 hours for LC-MS/MS analyses of 96 samples (9.4 min per sample). The tRNA modifications analytical platform was now applied to analyze the effects of 4,600 gene products on the levels of 41 tRNA modifications in *P. aeruginosa* PA14.

### tRNA modification profiling of a PA14 transposon insertion mutant library

To demonstrate the utility of the tRNA modification analytical platform, we used it to screen the non-redundant *P. aeruginosa* UCBPP-PA14 transposon insertion mutant library consisting of 5,746 mutant strains covering 4,600 non-essential genes^18^. This study required an initial identification of the tRNA modifications to be quantified and several foundational experiments to establish quality control parameters. To be as comprehensive as possible, we identified modifications reported in published studies of wild-type PA14^24, 25^ and modifications identified in a better-studied Gram-negative bacterium, *Escherichia coli*^26^ (**Supplementary Table 4**). Under our bacterial culture conditions, we identified 35 modifications that were detected in wild-type PA14 tRNA, with an additional 6 – cmnm^5^s^2^U, nm^5^s^2^U, mo^5^U, preQ_1_, oQ and s^2^U – that accumulate as modification pathway intermediates only in specific PA14 mutants. Out of 41 modifications, 37 were validated using synthetic standards. For the 4 modifications without synthetic standards, m^6^t^6^A and ms^2^io^6^A were tentatively identified by neutral loss analysis and confirmed by high-resolution mass spectrometry (**Supplementary Figure 10**)^27^. The remaining two modifications, ct^6^A and oQ, were identified by both quantifier and qualifier transitions, and their absence was noted in mutant strains lacking their synthetic enzymes (ΔTcdA and ΔQueA, respectively).

These 41 modifications were then used to evaluate the performance of the entire platform applied to 24 PA14 mutant strains in 4 biological replicates. For most of the RNA modifications analyzed, the coefficient of variation (CV) was below 25% (**Figure 1C**). However, ac^4^C^28^, ho^5^U^29^, m^1^A^30^, m^4^C^31^, and m^2,2^G^32^ all exhibited greater variation in the signal intensity and did not meet our reliability criteria. While these and the other modified ribonucleosides are all known to exist in bacteria and archaea^28–32^, the variability of detecting these four modifications was almost certainly due to their well-recognized chemical instability^33^, inherent low abundance, or possibly absence in PA14 (**Figure 1C**). To mitigate the risk of false positives in biological findings, these modifications were omitted from subsequent analysis. These studies defined a robust set of 30 tRNA modifications and established platform performance characteristics to have confidence in the results obtained with the 5,746-strain screen.

The validated platform was now applied to the complete 5,746-strain PA14 gene knockout library. The entire screen required ∼60 hours of Tecan robot time and ∼900 hours of LC-MS/MS time, generating >200,000 RNA modification quantifications. LC-MS/MS data were processed according to the flowchart depicted in **Supplementary Scheme 1**. The resulting fold-change data for 35 modifications in 5,746 strains are depicted in **Figure 2A**, where it is apparent that most of the modifications did not change by more than a 2-fold increase or decrease (gray points in **Figure 2A**). However, the loss of 312 genes caused levels of 30 modifications to change significantly (>2-fold; red and blue points in **Figure 2A**).

**Figure 2.**
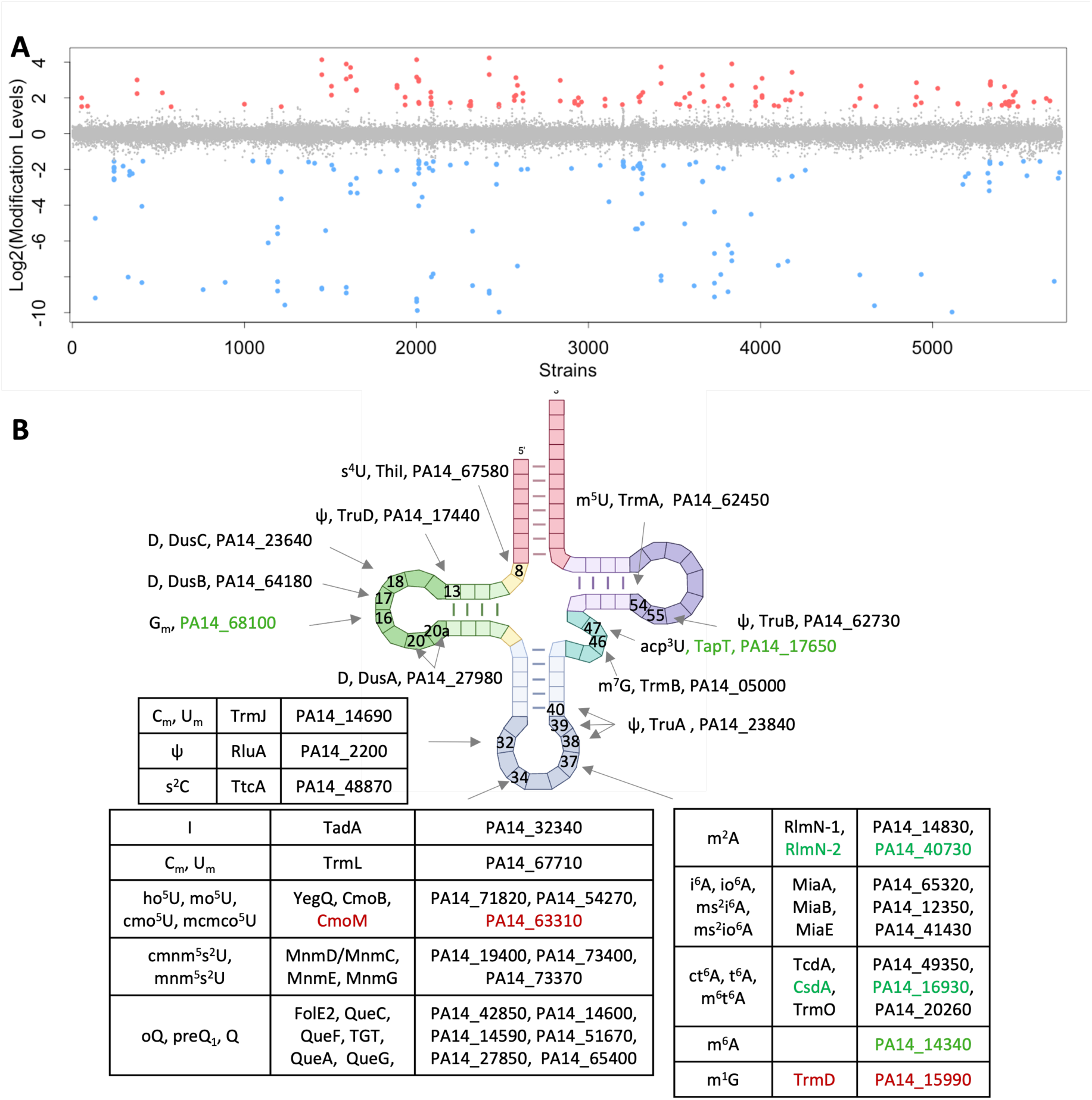
tRNA modification profiling in the PA14 mutant library reveals a constellation of gene-modification linkages. (**A**) Genes regulating the levels of tRNA modifications are visualized in this scatter plot of RNA modification fold-change values (calculated as described in Methods section, y-axis) across 5,746 strains in the PA14 gene knockout library (x-axis). Modifications with log_2_(fold-change) >1 are noted with red circles, <-1 in blue, and between log_2_(fold-change) ±1 in gray. (**B**) PA14 tRNA modifications and corresponding modifying enzymes validated and identified in this study. The unannotated proteins in green are newly identified as tRNA-modifying enzymes in this study. The mutants in red were listed in the library but sequencing and LC-MS analyses revealed that they were mislabeled.

As with any transposon insertion library, some care must be taken in interpreting the results. For example, loss of putative *uspA* (*PA14_41440*) increased io^6^A and the io^6^A/i^6^A ratio (**Supplementary Table 5**), suggesting that *miaE* expression or activity was upregulated (see **Figure 3D**). In the PA14 genome, *miaE* (*PA14_41430*) shares a promoter region with *uspA* in a ‘head-to-head’ orientation. One potential explanation for elevated MiaE levels in the *uspA* knockout is that accumulated stress in *uspA* mutant stimulates an alternative sigma factor to bind to the *uspA* promoter region, causing upregulation of *miaE* as a ‘side effect’. Another potential reason is that insertion of a transposon has obvious polar effects on the expression of downstream genes but can also affect the expression of upstream genes^34^. In this case, *miaE* is located ∼700 bp upstream of a transposon insertion site and its expression is upregulated. In two other cases, reduced tRNA modification levels are observed. For the *yajC* mutant, the insertion site was ∼200 bp downstream of the *tgt* gene, which could cause the observed reduction in Q (**Supplementary Table 5**), while the insertion site for the *thiG* mutant was ∼1000 bp upstream of the *trmB* gene, possibly causing the reduction of m^7^G (**Supplementary Table 5**). DNA sequencing analysis of several mutants was performed to confirm strain identities, revealing that several mutants differed from the library listings. The results have been summarized in **Supplementary Table 7**. For example, the transposon insertion site in the *trmD* (*PA14_15990*) mutant was located at an intergenic region, thus no reduction of m^1^G level was observed. As another example, repeated LC-MS/MS analyses of the two *aroB* knockouts PA14NR:38358 and PA14NR:42535 consistently showed cmo^5^U absent in PA14NR:38358 but present in PA14NR:42535. mRNA sequencing indicated that the regions upstream of the transposon insertion site are expressed at 10-fold higher levels than wild-type. For PA14NR:42535, the insertion site is only 32 bp from the translation initial site of *aroB* (**Supplementary Figure 11**). The resulting truncated *aroB* mRNA could produce a functional N-truncated AroB protein using an alternative translation start site, with 10-fold increased expression offsetting the reduced activity of the truncated AroB protein. A similar scenario might also be occurring in the mutant PA14NR:29841, where the transposon is inserted into the 5’ end of the *pdxA* gene, thus PdxA protein is still functional and no noticeable level change in corresponding RNA modifications as seen in another mutant PA14NR:40435 (**Supplementary Table 7**).

**Figure 3.**
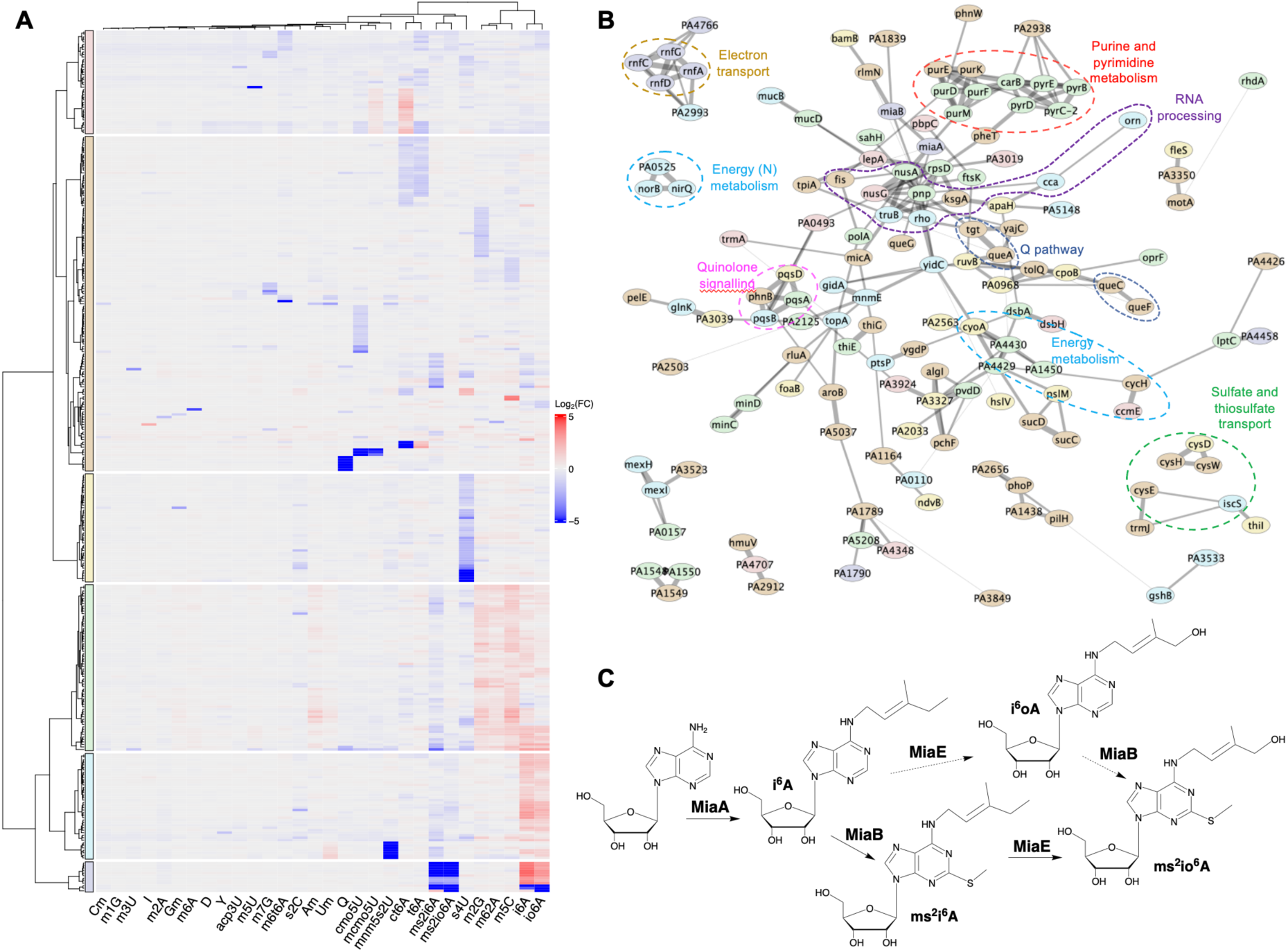
Visualization of gene networks regulating tRNA modification levels. (**A**) Hierarchical clustering of 351 PA14 mutants (including duplicates) with significantly altered levels of 30 tRNA modifications. The fold change calculation is described in Methods section. (**B**) String-database protein interaction network. The nodes (circles with gene names) represent 143 proteins in *Pseudomonas aeruginosa* PAO1 of which homologs in PA14 were encoded by genes that led to significant changes in the mutant library. The thickness of the edges (lines connecting two nodes) correlates with the StringDB confidence score of predicted protein interaction. The node colors correspond to the mutant clusters in the heatmap (right colored bars). For visualization, only clusters with ≥3 proteins and edges with confidence scores ≥ 0.5 are shown. (**C**) The biosynthetic pathways for i^6^A-derived tRNA modifications. Dashed lines represent proposed enzyme activities based on results from the PA14 screen.

Despite these few exceptions, the fidelity of most mutants appears to be high. For example, among the 351 mutants corresponding to 312 genes with altered tRNA modifications, 28 genes encode enzymes known to be involved in the synthesis of RNA modifications in PA14 (**Figure 2B**, **Supplementary Table 6**). The loss of these genes caused an absence or a decrease (for modifications modified by multiple writers) in the levels of specific RNA modifications (**Figure 2A**). While these confirmations further validate the platform, analysis of the full set of 312 genes causing modification changes revealed novel RNA modification genes and novel pathways linking environmental changes to changes in the tRNA epitranscriptome.

### Epitranscriptome pathway quantification defines substrate specificities

The dataset of 312 genes affecting 30 tRNA modifications was mined at several levels, starting with hierarchical clustering to identify co-variance among mutant strains (**Figure 3A**). In addition to the 28 known RNA-modifying enzymes, this analysis revealed clusters of genes associated with several groups of modifications, such as i^6^A and io^6^A, ms^2^i^6^A and ms^2^io^6^A, and several methylation modifications. Deeper analyses of gene-modification relationships revealed not only novel and unannotated gene function but also more subtle enzyme-substrate relationships *in vivo*. The synthesis of the i^6^A-related modifications illustrates the latter point. The levels of i^6^A and io^6^A, ms^2^i^6^A and ms^2^io^6^A are all reduced by loss of a single gene *PA14_65320*, *miaA*, the protein product of which is well established to catalyze the transfer of the isopentyl group from dimethylallyl diphosphate to the N^6^ position of A37 in tRNA to form i^6^A37 (**Figure 3C**), the precursor to the other three modifications. Conversely, i^6^A and io^6^A levels are negatively correlated with those of ms^2^i^6^A and ms^2^io^6^A when 49 genes are lost (**Figure 3A, Supplementary Table 5**). These results are consistent with known substrate/product relationships for the position 37 modifications and the 3 enzymes that catalyze their formation (MiaA, MiaB, MiaE). In particular, MiaE has been known to catalyze hydroxylation of ms^2^i^6^A to ms^2^io^6^A in other bacteria^35, 36^ (**Figure 3D**), but there is a claim that i^6^A is a poorer substrate for MiaE than ms^2^i^6^A in *Samonella typhimurium*^36^. Our results support the idea that MiaE prefers ms^2^i^6^A over i^6^A^37^: (1) the abundance for ms^2^io^6^A is ∼40-times higher than that of io^6^A and (2) the io^6^A/i^6^A ratio is ∼0.2, while the ms^2^io^6^A/ms^2^i^6^A ratio is ∼4 (**Supplementary Figure 12A**). Additionally, there is an observed increase in io^6^A levels by 6 to 9-fold in the two *miaB* mutant replicates in the library (**Supplementary Table 5**), which is consistent with io^6^A as a substrate for thiolation by MiaB to ms^2^io^6^A (**Figure 3C**).

### Discovering novel RNA modification activities

In addition to assigning *in vivo* substrate specificities, the 30-modification library screen provided functional annotation of RNA-modifying gene products and revealed evolutionarily distinct dual-function enzymes with non-redundant modification activities. The latter issue is important since LC-MS/MS analysis of RNA modifications in size-based small RNA fractions is susceptible to potential contamination with RNA fragments derived from mRNA, rRNA, and other long RNAs. This can lead to misidentification of modifications such as m^6^A and pseudouridine (4′) as tRNA modifications during analysis of the small RNA fraction. Our analytical platform offers a distinct advantage in providing corroborating evidence through the analysis of multiple RNA types from different mutant strains. For example, we identified PA14_14340 as a dual-specificity (adenosine-C6)-methyltransferase for both rRNA and tRNA. PA14 lacks YfiC, a known tRNA m^6^A writer that specifically modifies A37 of tRNA_1_^Val^ in *E. coli* ^38^. Instead, PA14_14340 is responsible for 99% of m^6^A modification in tRNA and 45% in rRNA, with the remaining 55% m^6^A in rRNA attributed to PA14_66340 (RlmJ) (**Supplementary Figure 13A,C**). PA14_14340 shares significant sequence and structural similarity with RlmF (**Supplementary Figure 14)**, an adenine-N6 methyltransferase specific for modification of A1618 of 23S rRNA in *E. coli* ^39^ (**Figure 4A, Supplementary Figure 13)**. In the docking model of PA14_14340 protein and tRNA^Val^, PA14_14340 formed a positively charged concave surface near the SAH-binding site surrounded by three polypeptide loops (boxed in **Figure 4A**) that likely interact extensively with the anticodon stem loop (ASL) from various angles like human METTL16^40^. The structural analyses support our findings that could function as a dual- specific m^6^A methyltransferase targeting both rRNA and tRNA.

**Figure 4.**
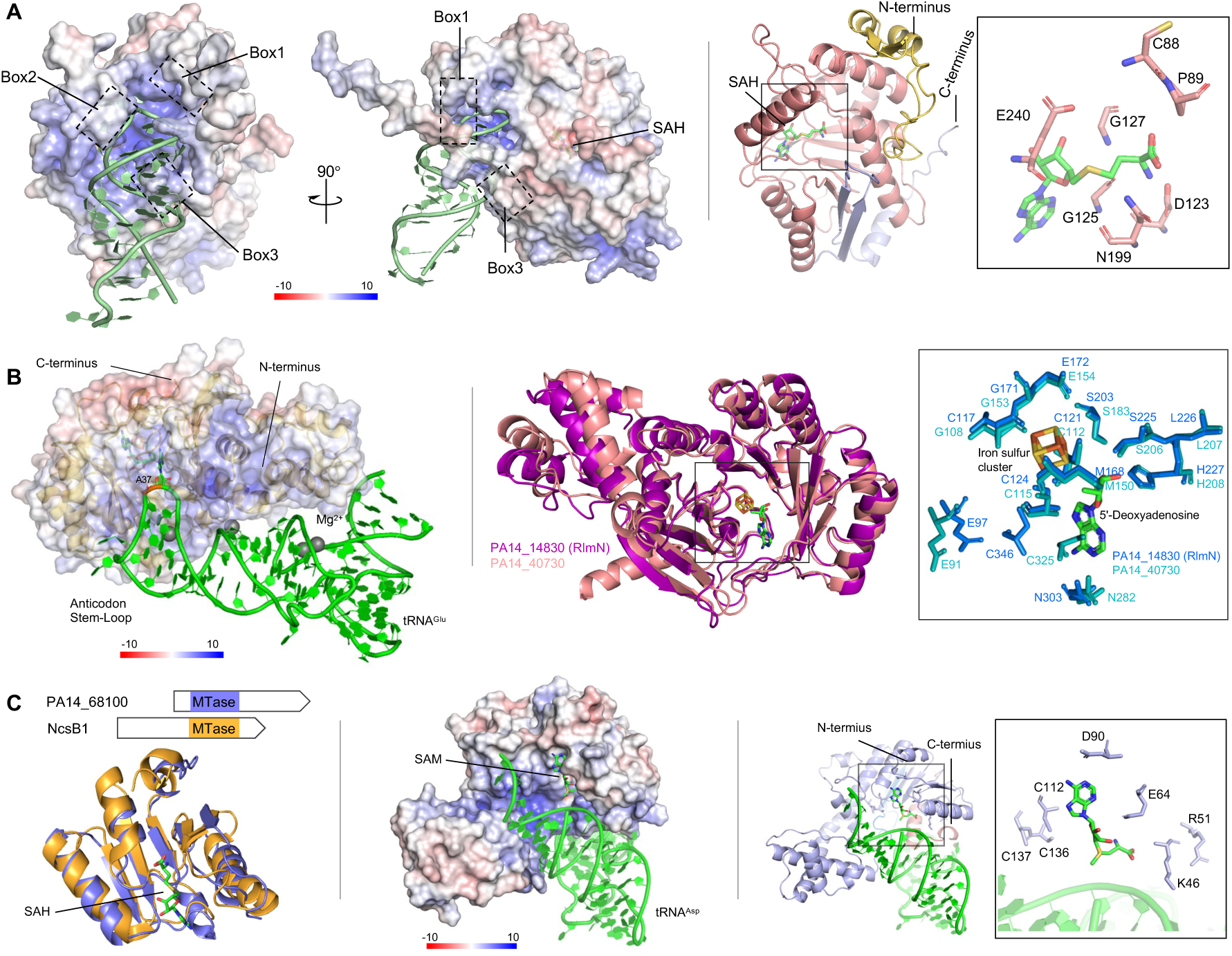
Structure modeling of PA14_14340, PA14_40730, and PA14_68100. (**A**) Left: Putative tRNA-binding site of PA14_14340. The anticodon stem loop (ASL) of tRNA^Val^ is represented by green cartoon. Three helix-turns that surround the groove of ASL are boxed. Right: the structure modeling of PA14_14340. The sticks boxed represent the key residues in the proximity of SAH. (**B**) Left: Putative tRNA-binding site of PA14_40730 protein. The tRNA^Glu^ is represented by green cartoon with A37 shown in stick format. Right: The structure alignment of PA14 RlmN and PA14_40730. The stick boxed represent the key residues in the catalytic pockets of the two proteins. (**C**) Left: structure alignment of the SAM dependent methyltransferase domain (MTase) of PA14_68100 and NscB1 (Uniprot Q84HC8). Middle: Putative tRNA-binding model of PA14_68100. The structure of tRNA^Asp^ is represented by light-green cartoon. Right: The structure modeling of PA14_68100. The sticks boxed represent the conserved residues in the proximity of SAH. Electrostatic potential mapped on the surface of each protein in which positive charges are shown in blue, negative charges in red, and neutral charges in white. SAH, [4Fe-4S]^2+^ cluster, 5′-dA, and SAM are shown in stick as indicated and colored by atom type.

Similarly, we identified two enzymes responsible for inserting m^2^A in tRNA. In *E. coli*, RlmN is the only m^2^A synthesis enzyme and modifies both rRNA and tRNA^41, 42^. We found that in PA14, the loss of *rlmN* (*PA14_14830*) eliminated m^2^A in rRNA and caused an 80% reduction in tRNA, while *PA14_40730* was responsible for the remaining 20% (**Supplementary Figure 13A,B**). Given that the abundance of m^2^A in tRNA is ten-times higher than in rRNA, the relatively small reduction in m^2^A in tRNA in the PA14_40730 knockout (**Supplementary Figure 13G**) is unlikely caused by rRNA contamination. Phylogenetic analyses showed that the RlmN branch comprises members from most bacterial taxa and its topology generally agrees with the universally accepted evolutionary history of bacteria (while the PA14_40370 branch includes representatives from a limited number of taxa; **Supplementary Figure 15**). This tree suggests that PA14_40730 evolved by duplication of the RlmN followed by a shift in substrate specificity. This is consistent with the significant sequence and structural similarity between PA14_40730 and RlmN. A positively charged groove in both proteins that can accommodate the anticodon stem-loop of tRNA of which A37 is placed in the active site^43^ (**Figures 4B, Supplementary 16**). The conserved residues Met168 and Cys346 (RlmN numbering) are likely involved in a transient thiosulfuranyl radical (boxed in **Figure 4B**), while amino acids unqiue to RlmN or PA14_40730, such as Arg198 in RlmN (boxed in **Supplementary Figure 16**), may help distinguish tRNA substrates^43, 44^, which await confirmation.

We also discovered that PA14_68100 represents a non-orthologous displacement of the *E. coli* tRNA guanosine-2’-O-methyltransferase TrmH^45^ and is responsible for nearly all G_m_ and about 25% of C_m_ modifications in tRNA, but not in rRNA (**Supplementary Figure 13A,D,E**). The G_m_ modification in rRNA is catalyzed by PA14_65190 (RlmB) (**Supplementary Figure 13A,D**). The observed reduction of G_m_ and C_m_ levels in tRNA in the *PA14_68110* mutant is likely an artefact caused by a transposon neighboring effect (**Figure 13A**), as noted earlier. PA14_68100 contains a methyltransferase domain (PF13679) near the N-terminus, forming a Rossmann fold-like structure with seven beta strands and six helix, similar to that found in 2,7- dihydroxy-5-methyl-1-naphthoate 7-O-methyltransferase, Ncsb1 (**Figure 4C**). A positively charged concave surface of the domain and conserved residues near SAM can accommodate an RNA substrate and SAM effectively (**Figures 4C, Supplementary Figure 17**). PA14_68100-like 2’-O-methyltransferase is primarily found in Pseudomonadota and has evolved differently from TrmH and RlmB (**Supplementary Figure 18**).

Finally, we functionally confirmed tRNA modifying activities in PA14. For instance, loss of *PA14_16930* abolished ct^6^A (**Supplementary Table 5, Supplementary Figure 19B**). PA14_16930 shares strong sequence and structural similarity with *E. coli* cysteine desulfurase CsdA, which Suzuki and coworkers showed is one of three Csd family proteins required for ct^6^A formation^46, 47^ (**Supplementary Figure 19D**). Similarly, the deletion of *PA14_17650* resulted in an approximate 70% reduction of acp^3^U, mirroring the function of *E. coli* TapT (YfiP) as a tRNA-uridine aminocarboxypropyltransferase in PA14. This functional similarity is supported by sequence and structural alignments of PA14_17650 and TapT (**Supplementary Figure 20**), reinforcing recent findings in PA14^48–50^.

### Epitranscriptome regulatory networks

The knockout library epitranscriptome dataset provides an opportunity to identify gene networks that indirectly regulate specific tRNA modifications and families of related modifications, as illustrated by the six clusters in the heat map in **Figure 3A**. These networks also suggest possible regulatory roles for the modifications. At the first step away from RNA- modifying enzymes, multiple cofactor metabolism pathways were found to have a direct impact on RNA modification levels. For instance, loss of the *sahH* gene (*PA14_05620*), which encodes *S*-adenosyl-L-homocysteine (SAH) hydrolase, caused a decrease in SAM- dependent modifications such as U_m_, C_m_, m^5^U, m^3^U, m^7^G, ms^2^i^6^A, ms^2^io^6^A, mnm^5^s^2^U, cmo^5^U, mcmo^5^U, Q, but an accumulation of preQ_1_ (**Figure 5; Supplementary Table 5**). Loss of SahH activity leads to an accumulation of SAH as the byproduct of methyltransferase cofactor *S*- adenosyl-L-methionine (SAM). SAH is known to be a non-selective feedback inhibitor for many methyltransferases^51^, which explains the reduced levels of methylation-based modifications.

**Figure 5.**
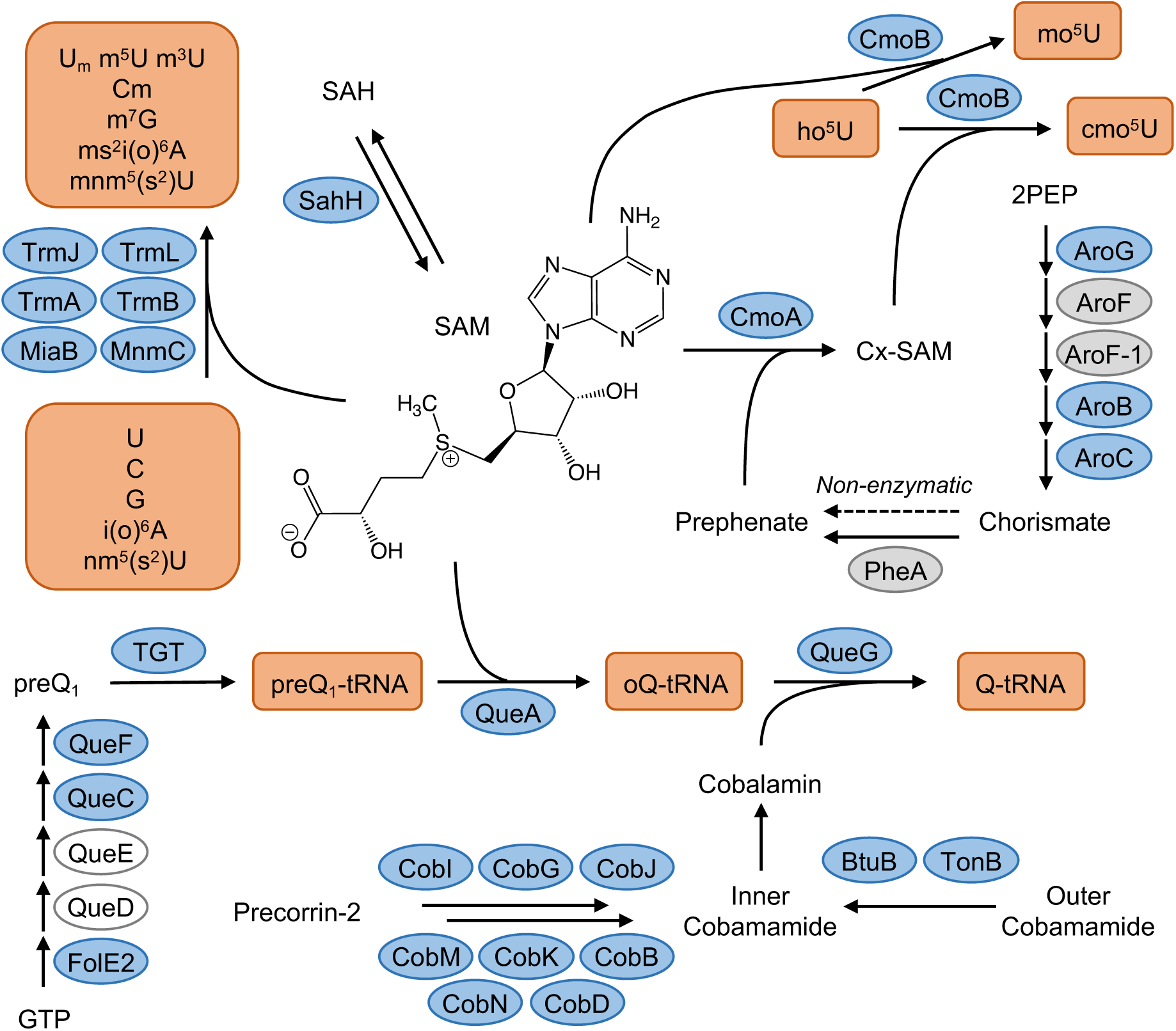
Genome-scale analysis of tRNA modifications in PA14 mutant library reveals a SAM-centric gene network influencing tRNA modification biogenesis. SAM and SAM analogue carboxyl-SAM (Cx-SAM) participate in methylation of tRNA modifications with a variety of methyltransferases. For example, CmoB differentially catalyzes mo^5^U and cmo^5^U depending upon the availability of SAM and Cx-SAM, respectively, with Cx-SAM levels determined by levels of prephenate from the shikimate pathway. SAM is also involved in the biogenesis of Q, together with multiple tRNA modifying proteins and the QueG cofactor cobalamin. Proteins in blue ellipses significantly affected levels of tRNA modifications noted in the orange boxes. Proteins in grey ellipses do not affect tRNA modification levels. Clear ellipses represent proteins for which the encoding genes are absent in the PA14 knockout library. The dashed arrow linking chorismate to prephenate denotes a non-enzymatic conversion and the solid arrow denotes chorismate’s involvement in ho^5^U formation/ PEP: 2-phophoenolpyruvate.

Another example involves carboxy-S-adenosylmethionine (Cx-SAM), one of the cofactors for CmoB-mediated synthesis of cmo^5^U and mcmo^5^U (**Figure 5**). Loss of *aroB* and *aroC* in the shikimate pathway, involved in aromatic amino acid synthesis, reduced cmo^5^U and increased ho^5^U and mo^5^U (**Figure 3A, 5; Supplementary Table 5**). The shikimate pathway produces prephenate, a substrate for CmoA synthesis of Cx-SAM (**Figure 5**). CmoB preferably uses Cx-SAM over SAM to produce cmo^5^U^52^. In the absence of Cx-SAM, CmoB can still use SAM to synthesize mo^5^U, albeit at a reduced rate. Loss of *aroB* and *aroC* fully blocks the synthesis of prephenate and Cx-SAM, thus leading to the disappearance of cmo^5^U and accumulation of mo^5^U. The loss of *pheA* in this pathway (**Figure 5**) did not completely eliminate cmo^5^U, which is consistent with the slow non-enzymatic conversion of chorismate to prephenate^53^. Additionally, at least 3 redundant 3-deoxy-D-arabinoheptulosonate 7-phosphate (DAHP) synthases were identified in PA14 genome: AroF, AroF-1, and a putative AroG (PA14_27330) (**Figure 5**). Deletion any one of these enzymes had no impact on mo^5^U or cmo^5^U levels (**Supplementary Table 5**).

Expanding the modification network outward, deletion of genes involved in cobalamin translocation (*btuB, tonB*) and biogenesis (*cysG, cobB/D/G/I/J/K/M/N/H*) all impeded the cobalamin-dependent QueG conversion of epoxyqueuosine (oQ) to Q (**Figure 5**) and accumulation of oQ (**Supplementary Table 5**)^54^. Interestingly, loss of the anaerobic cobalamin biogenesis enzyme CysG led to the accumulation of oQ even under aerobic conditions, suggesting that the CobA aerobic counterpart might not fully compensate for CysG function.

A broader perspective on gene networks affecting tRNA modification levels is achieved by translating the hierarchical clustering in **Figure 3A** to a network map such as that shown in **Figure 3B**. Here we used the STRING protein interaction database^55^ to evaluate interactions among 143 proteins (nodes/circles with gene names) the loss of which significantly altered individual tRNA modifications. In several instances, the six clusters of mutant strains in the heat map (**Figure 3A**) translate to clusters of proteins in the network map (**Figure 3B**; node colors = heat map clusters), which reflects one level of functional relatedness of proteins affecting tRNA modifications. This functional relatedness is emphasized when Gene Ontology (GO) categories are overlaid on the protein interaction network (dashed circles in **Figure 3B**), which suggests a potential regulatory role for tRNA modifications in various aspects of bacterial physiology. This regulatory potential is apparent in another form of network analysis in which each modification is linked to a gene mutation that affects the modification level, as shown in **Supplementary Figure 12B**. For example, i^6^A was altered by 61 genes (nodes) and connected with io^6^A through another 35 genes, more than any other pair of modifications. The pairs ct^6^A-t^6^A and m^6^t^6^A-ms^2^io^6^A were connected by 17 and 10 nodes, respectively, as they share the same synthesis pathways. As a node, *PA14_05620* (*sahH*) connects to the most modifications (11), which is consistent with its role in recycling the SAH product of the SAM cofactor for RNA methyltransferases in PA14 as noted earlier.

The regulatory potential for the tRNA epitranscriptome is also apparent in connections between individual modifications. For example, 35 of 39 nodes in connected with ms^2^i^6^A in **Supplementary Figure 12B** are also connected with other modifications, including i^6^A, m^6^t^6^A, ms^2^io^6^A, m^5^C, m^3^U, and m^2^G. This suggests a role for ms^2^i^6^A as, for example, a precursor in tRNA maturation that affects installation of other modifications or as part of a signaling network linking metabolic shifts to translation. This latter point is illustrated next with the i^6^A family of tRNA modifications.

### The MiaB Fe-S cluster as an integrator of metabolic signaling

The levels of the i^6^A family of modifications (i^6^A, io^6^A, ms^2^i^6^A, ms^2^io^6^A) are affected by dozens of genes in PA14 (**Figure 3A, Supplementary Figure 12**), which raises the possibility that the enzymes catalyzing these modifications (MiaABE in **Figure 3C**) function as sensors in signaling pathways with regulatory links to translation. Here we explored this idea with MiaB, which inserts a sulfur in i^6^A to form ms^2^i^6^A and possible into io^6^ to form ms^2^io^6^A^56^, by analyzing the PA14 genes that caused the ms^2^i^6^A/i^6^A ratio to fall below 0.5, which indicates inhibition of MiaB activity. A map of protein-protein interactions for these genes (**Figure 6A**) reveals a complicated but significant (PPI enrichment *p* <1.0e^-16^) network of 104 gene nodes with 132 edges, reflecting cellular processes such as iron-sulfur cluster assembly and maintenance, nitric oxide detoxification, and oxidative stress response. It is immediately apparent that the MiaB Fe-S cluster plays a key role in this network, as indicated by the strong association with Fe-S cluster biogenesis and maintenance proteins: Cysteine desulfurase IscS and the PdxA/J enzymes involved in the synthesis of its cofactor pyridoxal 5’-phosphate (PLP)^57^; BfrB for iron storage^58^; the RnfA/C/D/G/H family as the ferredoxin/electron donor^59^; GshA/B for Fe-S cluster export^60^; and GrxD for the maintenance and repair of Fe-S cluster proteins^61^. Loss of these proteins inhibited the conversion of i^6^A to ms^2^i^6^A, especially for proteins in the Rnf complex, loss of which caused the complete disappearance of ms^2^i^6^A.

**Figure 6.**
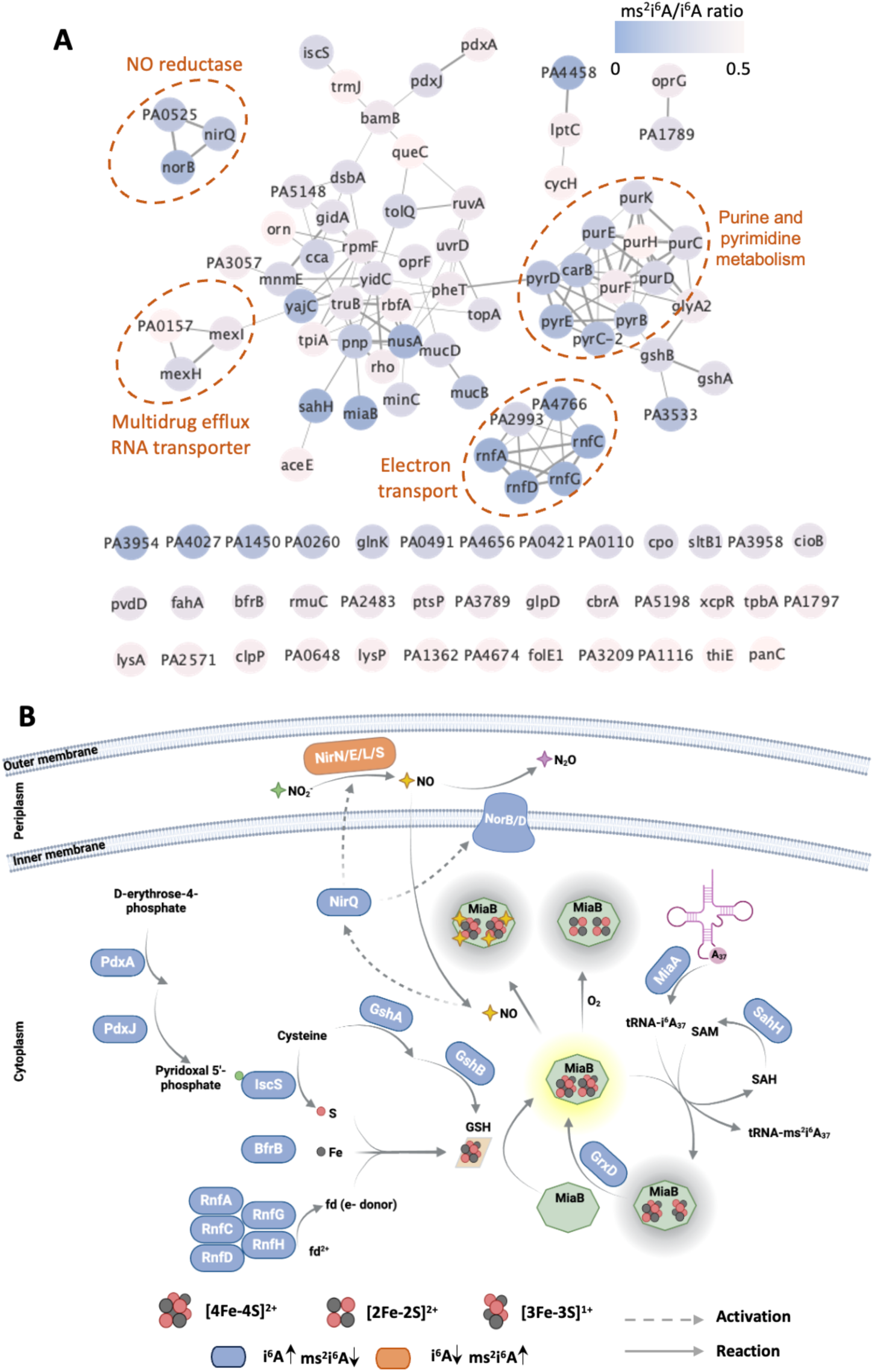
Visualization of gene networks influencing MiaB activity. (**A**) Here we used the ratio of ms^2^i^6^A to i^6^A (ms^2^i^6^A/i^6^A < 0.5 ) as metric to create an interaction network of proteins affecting MiaB activity. The genes are annotated according to *Pseudomonas aeruginosa* PAO1 strain. Node color indicates the ms^2^i^6^A/i^6^A ratio (key upper right), the edge width (connecting lines) correlates with the stringDB confidence score for the protein interactions; confidence scores ≥ 0.4 are displayed. Functional categories are encircled with a dashed line. (**B**) Diagram illustrating the metabolic and regulatory pathways centered on Fe-S clusters affecting MiaB’s activity and ms^2^i^6^A biosynthesis, as deduced from Figure 5A. Proteins highlighted in blue represent gene knockouts that lead to MiaB dysfunction, increasing i^6^A levels while decreasing ms^2^i^6^A. Conversely, proteins in orange represent gene knockouts that lower NO levels, thereby enhancing MiaB activity, resulting in decreased i^6^A and increased ms^2^i^6^A levels.

The signaling network expands when other gene clusters in **Figure 6A** are considered. NO metabolism regulates a variety of pathogenesis pathways in the facultative anaerobe *P. aeruginosa* at low oxygen tensions (denitrification alternative respiration), as well as from NO exposure from environmental exposures such as activated macrophages, by activating the dissimilative nitrate respiration regulator (DNR) transcription factor^62^. Genes *nirN*, *nirE*, *nirL*, *nirS*, *nirQ*, *norB* and *norD* are involved in nitric oxide (NO) metabolism. Loss of *nirQ*/*norB*/*norD* leads to accumulation of NO, which is well established to be disrupt Fe-S clusters by forming a Fe-S-NO complex^63^. Conversely, with the deletion of the nitrite reductase genes *nirN*, *nirE*, *nirL*, and *nirS*, an increase in MiaB activity was inferred from the observed reduction of i^6^A and io^6^A levels, coupled with a slight rise in ms^2^i^6^A levels (**Supplementary Table 5**). Another similar and perhaps related gene cluster involves oxidative stress and reactive oxygen species (ROS) response proteins: glutathione biosynthesis enzyme GshA/B^64^, redox enzyme glutaredoxin GrxD^65^, vitamin B6 synthesis enzymes PdxA/J^66^, alkanesulfonate monooxygenase SsuD (PA14_12710)^67^, pyrimidine biosynthesis (PyrC/D/E, PA14_05250/24640/70370)^68, 69^, sigma factor algU regulator MucB (PA14_54410)^70^ and DNA- binding transcriptional regulator OxyR (PA14_06400)^71^. In parallel, ROS levels increase with loss of proteins for transcription termination factor NusA (PA14_62770)^72^, transcription termination factor Rho (PA14_69190)^73^, multidrug efflux RND transporter MexH/I (PA14_09520, PA14_09530)^74^, cmnm^5^s^2^U writer MnmE/G (PA14_73400, PA14_73370)^75^, and oligoribonuclease Orn (PA14_65410)^76^.

## Discussion

RNA modifications have been studied for decades, but only recently has the development and application of ‘omic technologies led to the discovery of their systems-level function in regulating gene expression at the level of translation^5, 10, 77–79^. One of the physiological functions of the system of tRNA epitranscriptome reprogramming and codon-biased translation is adaptation to environmental changes and stress^5, 77, 78, 80^. However, the gene and signaling networks regulating the levels of tRNAs and tRNA modifications in this system have not been defined, in part due to the lack of high-throughput technology for large-scale functional genomics studies. Toward the goal of understanding epitranscriptome regulatory networks, we developed a high-throughput tRNA modification quantification platform and applied it to 5850-strain *P. aeruginosa* PA14 transposon insertion library. The results provide the first comprehensive dissection of tRNA modification regulatory networks, while the flexibility of the platform opens the door to sequencing- and LC-MS-based ribonucleomics and epitranscriptome mapping in any type of biological sample.

Driven by RNA instability^81^ and small sample size, the quantitative precision (CV <25%) and accuracy that we were able to achieve with this high-throughput epitranscriptome analysis platform required optimization of the five steps of cell lysis, RNA purification, RNA processing, MS analysis, and MS data processing. We identified several critical parameters that affected precision and accuracy: (1) a lysis buffer with a high concentration of guanidine thiocyanate more efficiently releases RNA, rapidly inhibits RNases and RNA-modifying enzymes, and enhances the size-based resolution of the magnetic beads (**Supplementary Figure 1**); (2) the components of the large RNA binding buffer I determined the efficiency of removal of large RNA without affecting tRNA (**Supplementary Figure 2**); (3) isopropanol concentration during the tRNA binding step affected the yield of tRNA and co-precipitation of other nucleic acid metabolites (e.g., ATP); (4) carboxyl magnetic beads were chosen because of their compatibility with all different sample types, including bacteria, mammalian cells, and tissues (**Supplementary Figure 3**); and (5) workflows and the choice of plasticware for automated tRNA isolation using a liquid handler affected the reproducibility and purity of the extracted tRNA (**Supplementary Figures 4-6**). Care must be taken in interpreting RNA modification data derived from size-based RNA purification given the modifications present in small biologically relevant RNA fragments from rRNA, mRNA and other long RNAs. Our focus on tRNA modifications reduces the abundance of non-tRNA contributions given the much higher density of modified ribonucleosides in tRNA compared to other forms of RNA^82^, though we are cautious in interpreting quantitative data for modifications that appear in both tRNA and rRNA (e.g., Gm, Cm, Um, Am). Care must also be taken to avoid artefacts arising from DNA processing. For example, enzymatic hydrolysis of RNA is prone to deamination of C, G, and A by contaminating deaminase enzymes, which necessitates the use of deaminase inhibitors^83^. Similarly, antioxidants must be added to prevent loss of oxidation-sensitive modifications such as ho^5^U and ho^5^C^84^. We also observed a bias toward nonpolar RNA modifications when using 10,000 Da MW cut-off centrifugal filtration devices to remove enzymes (**Supplementary Figure 9**).^85^ While use of a 2 µm inline filter positioned between the injector and the LC column protects against particulate contamination, the enzymes still enter the LC column and risk performance degradation. However, such degradation was not apparent in our analysis. Here we developed a rapid UPLC-MS/MS method that minimizes analysis time and maximizes ribonucleoside resolution in a 6-minute run for 40 modifications. This amounted to ∼900 h for 5,850-strain library. Clearly, the platform can be adapted to focus on any modifications in any organism or tissue for sensitive quantification of RNA modifications from any form of RNA.

In addition to enhancing functional annotation of RNA modification-related genes, application of the epitranscriptome platform revealed several tRNA modification gene networks that have significant regulatory potential. Among the most dynamic modifications in this analysis, i^6^A, io^6^A, ms^2^i^6^A, ms^2^io^6^A, Q, cmo^5^U, mcmo^5^U, and ho^5^U are located the wobble position of the anticodon or at position 37, both of which play central roles in regulating codon recognition. Importantly, a wide variety of stresses have been shown to regulate tRNA modifications and cause selective translation of codon-biased mRNAs encoding stress response proteins^5, 8, 9, 80, 86–88^. Screening the PA14 knockout library for changes in tRNA modification levels provides new insights into the mechanisms linking the tRNA epitranscriptome to both stress sensors and stress response effectors. The dynamics of the xo^5^U wobble modification family illustrate this point. Coupled with the observation that the chorismate precursor to prephenate regulates ho^5^U formation^89^ (**Figure 5**), reductions in cmo^5^U and mcmo^5^U and increases in ho^5^U and mo^5^U all caused by loss prephenate synthetic genes AroBCG point to a potential regulatory cycle. In this scenario, cellular decreases in prephenate, due to increased demand for aromatic amino acids or secondary metabolites from the shikimate pathway leads to reduced levels of Cx-SAM, reduced levels of cmo^5^U and mcmo^5^U modifications in tRNAs with UNN anticodons: tRNA-Ala-<u>U</u>GC, tRNA-Ser-<u>U</u>GA, tRNA-Pro-<u>U</u>GG, and tRNA-Thr-<u>U</u>GU for mcmo^5^U and tRNA-Leu-<u>U</u>AG and tRNA-Val-<u>U</u>AC, for cmo^5^U^90^. Since cmo^5^U and mcmo^5^U expand the codon repertoires of these tRNAs to include G-ending codons^91, 92^, we would predict that reduced levels of these modifications would shift translation to favor A- and T- ending codons.

The dynamics of MiaB-mediated i^6^A modifications further illustrates the translational regulatory potential of tRNA-modifying enzymes. Just as there are Fe-S cluster-containing transcriptional regulators that sense O_2_ and iron^93, 94^, it is reasonable to propose Fe-S cluster enzymes as translational regulators given their widespread role in the formation of RNA modifications^95^. MiaB has the potential to serve as such a translational regulatory node given its heightened sensitivity to gene knockouts in the Fe-S biogenesis, repair, and redox regulation pathways. Other Fe-S cluster RNA-modifying proteins, such as QueG (Q), TtcA (s^2^C), and RlmN (m^2^A), were not affected by the loss of genes that significantly altered MiaB activity. For example, deletion of *nirQ, norB,* or *norD* did not cause noticeable changes in Q, s^2^C and m^2^A levels but did alter i^6^A-related modifications significantly (**Supplementary Table 5**). One potential reason is that MiaB is unusual in containing two [4Fe-4S] clusters, one of which mediates formation of the adenosine radical intermediate and the other forms an unstable [3Fe-4S] intermediate involved in the methylthio group transfer reaction^96, 97^. The resulting [3Fe-3S] cluster must be repaired to restore MiaB activity. The Fe-S clusters of QueG, TtcA, and RlmN, on the other hand, remain intact during transfer of an electron transfer, sulfur, or methyl group, respectively^54, 98, 99^. With i^6^A located specifically in the subset of tRNAs that read UNN codons^100^, the dependence of MiaB activity on Fe-S biogenesis, repair, and redox regulation pathways suggests that i^6^A and ms^2^i^6^A dynamics could reprogram the tRNA pool to regulate translation of stress response mRNAs based on their UNN codon content, as proposed for the xo^5^U modifications.

While our method has yielded promising results, there are still several limitations that future advancements in this technology could address for broader applications. Like any other existing tRNA purification methods, the method here is not able to provide 100% pure tRNA, so the results are somehow biased by tiny rRNA traces. For example, the m^4^C and m^6^_2_A are supposed to be rRNA modifications. The presence of m^5^C in bacterial tRNA is under debated and its modifying enzyme is not resolved yet, which needs m^5^C mapping method to further validate the results. Additionally, the RNA hydrolysis method employed in this study may not be ideally suited for the analysis of labile modifications, such as ct^6^A and t^6^A. ct^6^A hydrolyzes to t^6^A under basic conditions and these modifications are prone to Tris adduct formation and alkaline-induced epimerization (**Supplementary Figure 19A**)^101^. While this potential source of bias did not impede the identification of associated proteins, such as ct^6^A-forming CsdA and CsdL (**Supplementary Table 5**), refining the RNA hydrolysis protocol will be important for studies specifically targeting these two modifications to ensure analytical accuracy and precision. One feature of the method that will need improvement involves reducing the input sample size to accommodate adherent mammalian cells cultured in 96-well plates or limited quantities of tissue. In this study, we used ∼0.3 OD_600_ of PA14 to extract ∼1 μg of tRNA. However, for mammalian cells grown in a 96-well plate, we anticipate obtaining ∼60 ng of tRNA from ∼30,000 cells. This smaller quantity of tRNA requires refinement of the liquid handler plasticware and workflow to manage smaller working volumes, as well as adjustments to the LC-MS/MS system for increased sensitivity. Another important improvement would be to shorten the turnaround time for each LC-MS/MS run, especially when conducting larger scale screening. Finally, when dealing with the large mass spectrometer datasets, AI-enabled tools would be very useful, such as supervised machine learning for efficient peak detection to minimize the need for manual curation, as well as biomarker detection and gene network analysis.

In summary, we developed a robust and high-throughput approach to quantitative analysis of the tRNA and rRNA epitranscriptomes. These results reveal the impact of the tRNA analytical platform on defining the layers of gene-gene interactions affecting tRNA modifications, expanding from catalytic genes to the larger circle of genes regulating cofactor production and further informing on gene-gene interactions at the level of signaling networks. The platform piloted with a gene knockout library can be readily adapted to large collections of human cells and tissues, such as the 2000 cancer cell types in the DepMap^15^ ^102^. The platform can also be adapted to other RNA analytical methods, such as small RNA processing for NGS library preparation for sequencing-based modification maps and quantification of small RNAs^103^. The 96-well plate-based RNA extraction and analysis capabilities also position the platform for drug discovery with both whole cell phenotypic and target-based screening of compound libraries, as well as complex biomarker discovery.

## Methods

### Cell culture and lysis

The 5,764-strain non-redundant library of *Pseudomonas aeruginosa* PA14 transposon insertion mutants^18^ was obtained from Dr. Deborah Hung at the Massachusetts General Hospital. Cell culture conditions followed the instructions provided in “The PA14 Non-Redundant Set of *Pseudomonas aeruginosa* Transposon Insertion Mutants User Manual (version 2.2)”. A 96-pin replicator was used to inoculate from frozen culture stocks to 0.3 mL LB medium containing either gentamycin (15 µg/mL) or kanamycin (200 µg/mL) in a 1.2 mL low profile 96-well plate (BRAND, 701340). The plate was sealed using a breathable film (Sigma, Z763724) and shaken overnight at 37 °C, 300 rpm. Then 100 µL of overnight culture was transferred to a 2.0 mL deep 96-well plate (VWR, 76329-998) containing 700 µL of LB medium and either gentamycin (15 µg/mL) or kanamycin (200 µg/mL). The plate was sealed and shaking continued at 37 °C, 300 rpm, until the cell density reached ∼0.8 OD_600_ (∼1.6 x 10^8^ CFU/mL). Cells were then pelleted by centrifugation of 3000 xg for 10 min at 4 °C. Medium was removed and cells were washed with cold 1× PBS buffer. After re-pelleting the cells, the PBS was discarded. Cell lysis buffer (100 µL; 50 mM Tris-HCl, 1 mM EDTA, 4 M GITC, pH 7.5) was added to each well and the plate was shaken vigorously at ambient temperature (1500 rpm) for 10 min to ensure complete cell lysis. The quantity of cell lysate was sufficient for 2 separate RNA isolations.

### RNA purification

Large RNA binding was performed by mixing 45 µL of cell lysate with 105 µL of RNA binding buffer I containing 3 M LiCl, 7% PEG8000, 2 mM EDTA, 40 mM Tris buffer (pH 7.5), and 8 µL Sera-Mag carboxylate-modified magnetic beads (Cytiva, 65152105050350). Under these conditions, large RNAs, but not small RNAs, bind to the carboxylate-coated magnetic beads. The mixture was agitated thoroughly and incubated at ambient temperature for 5 min. The magnetic beads were separated on a magnetic rack at ambient temperature for 5 min and the supernatant was transferred to a new tube/96-well plate. The tube/plate was then centrifuged at 3000 xg for 1 min and placed on the magnetic rack for 5 min to remove any remaining beads. The supernatant was transferred to a new tube/plate and mixed with 1.8 x (v/v) RNA-binding buffer II (0.05% magnetic beads in isopropanol). The mixture was agitated thoroughly and incubated at ambient temperature for 5 min. The magnetic beads were separated on the magnetic rack for 5 min and the supernatant was removed. The pelleted beads from 1^st^ round and 2^nd^ round binding reactions were washed twice using washing buffer (10 mM Tris-HCl, 80% EtOH, pH 7.5). The supernatant was removed and the beads were air dried at ambient temperature. Large RNAs were eluted in 50 µL nuclease-free water at 90 °C for 2 min. Small RNAs were eluted in 50 µL nuclease-free water at ambient temperature.

### Robotic workflow for RNA isolation

A robotic liquid handler (Tecan EV150, Switzerland) was used to complete the magnetic beads-based RNA isolation in 96-well plate format. The layout of the worktable is shown in **Supplementary Figure S3**. *<u>Large RNA binding</u>*: The plated cell lysates were placed on ice to thaw while reagents were prepared in 96-well plates for RNA isolation. The plate containing thawed cell lysates (45 μL) was placed in deck position 1. A 96- chanel pipette (MCA96) was used to transfer cell lysates to plate containing RNA binding buffer-1 (105 μL, deck position 5), and mix the reagents with 5 mix cycles. The plate then was left for sufficient rRNA and magnetic beads binding. *<u>Wash buffer aliquot</u>*: During this period, the wash buffer was being aliquoted (130 μL ×5) from trough “Wash Buffer” to plate “LRNA WB” (deck position 4). *<u>Beads pulldown (1)</u>:* The plate then was transferred by the robotic manipulator arm (RoMa) to magnetic plate (Alpaqua, Catalyst 96, A000550) “Magnet1” (deck position 10) for magnetic beads pulldown. *<u>Wash buffer aliquot:</u>* During the awaiting time, the wash buffer was being aliquoted (130 μL× 5) from trough “Wash Buffer” to plate “tRNA WB” (deck position 12). *<u>Beads pulldown (2)</u>:* The supernatant (145 μL) was then transferred to plate “Supernatant1” (deck position 6), and the residue was removed and discarded in waste plate “LRNA Waste” at deck position 8. Upon completion, the program was suspended, the plate “Supernatant1” was manually removed and placed in a centrifuge (3000 xg, 1 min) to spin down remaining 1^st^ round beads. The plate was placed back on magnetic plate “Magnet2” (deck position 9) and the program continued. *<u>Beads wash</u>:* During the waiting time for the 1^st^ round beads residue pulldown on "Magnet 2”, the 1^st^ round beads in plate “LRNA MB” on “Magnet1” (deck position 10) were washed with an aliquot of wash buffer from “LRNA WB” (300 μL). The wash buffer residue was removed and discarded after 60 s. After repeating the wash step, the 1^st^ round beads were left to air dry. *<u>tRNA binding</u>:* During this period, supernatant (120 μL) in plate “Supernant1” on “Magnet2” was transferred to plate “tRNA MB” (deck position 7) and mixed thoroughly with RNA binding buffer II (228 μL). *<u>Large RNA elution</u>:* While the tRNA was binding to the magnetic beads, nuclease-free water (55 μL) in a trough was added to the air-dried 1^st^ round beads in plate “LRNA MB”. The plate was manually removed and shaken on an orbital plate shaker (1000 rpm, 60 s), then incubated in a metal beads bath (90 °C, 90 s). Then the plate was placed back on “Magnet2” for 20 s for pulldown of the beads. The elutes (50 μL) were transferred to plate “LRNA Elute” (deck position 2), the plate was placed on ice immediately. *<u>Beads pulldown</u>:* The plate “tRNA MB” was transferred to “Magnet1” to pull down the beads. The supernatant was removed through three separate dispensing actions, with a 60 s interval between each to allow sufficient time for the magnetic beads to fully settle. This stepwise approach is essential due to the significant distance between the magnet plate and the beads, preventing incomplete bead recovery in a single dispensing action with substantial bead loss. *<u>Beads wash</u>:* The 2^nd^ round beads were washed by aliquoted wash buffer from “tRNA WB” (300 μL). The wash buffer residue was removed and discarded after 60 s, then the 2^nd^ round beads were left air dry. While awaiting the beads airdry, the preparation for next run can be performed. *<u>tRNA elution</u>:* Nuclease-free water (55 μL) in trough was added to airdried 2^nd^ round beads in plate “tRNA MB” on “Magnet1”. The plate was manually taken out and shaken on an orbital plate shaker (1000 rpm, 60 s). The plate was put back on “Magnet1”, wait 20 s for beads pulldown. Upon the elutes (50 μL) were transferred to plate “tRNA Elute” (deck position 3), the plate was kept on ice immediately until transferred for LC-MS sample preparation or stored at -80 °C for long-term storage.

### RNA hydrolysis for LC-MS/MS analysis

The purified RNAs in the 96-well PCR plate (Axygen Scientific, PCR-96-FS-C) were transferred to a 0.3 mL 96-well plate (Agilent, 5043-9313) for hydrolysis in a 40 µL of an enzyme cocktail containing 10 U benzonase (Sigma, 34998-4L), 4 U calf intestinal alkaline phosphatase (Sigma, 524572), 0.1 U phosphodiesterase I (US Biological, P4072), 0.1 mM deferoxamine (Sigma, D9533-1G), 0.1 mM butylated hydroxytoluene (Sigma, cat. #W218405), 4 ng coformycin (NCI, 27781713), 50 nM internal standard [^15^N]5-deoxyadenosine, 2.5 mM MgCl_2_, and 5 mM Tris-HCl buffer pH 8.0. The reaction mixture was incubated at 37 °C for 6 h.

### Liquid chromatography-coupled tandem mass spectrometry

#### Dynamic MRM scans

For RNA modification retention time validation (**Table S2**), synthetic standards were utilized in tandem with a Waters ACQUITY UPLC BEH C18 column (50× 2.1 mm, 1.7 µm) equipped with an 0.2 µm stainless steel inline filter (Waters, 205000343) connected to an Agilent 1290 Infinity II UHPLC system and an Agilent 6495 triple-quadrupole mass spectrometer. The LC was operated at 25 °C with a flow rate of 0.35 mL/min. Initial conditions held 100% solution A (water, 0.02% formic acid) for 2 min, followed by 2-4 min at 0-8% solution B (70% acetonitrile, 0.02% formic acid), and from 4-5.9 min at 8-100% solution B. The mass spectrometer was operated in positive mode with an electrospray ionization source with following parameters: gas temperature of 200 °C, gas flow of 11 L/min, and a capillary voltage of 3000 V. Detection leveraged a dynamic MRM mode, targeting product ions from precursor ions of each RNA modification. The collision energy (CE) was optimized by MassHunter Optimizer for maximal sensitivity for the modification. The hydrolysed RNAs in the 96-well plate were sealed with easy piercing film (BioChromato, REPS001, Japan), and kept at 5 °C during analysis.

#### Neutral loss scan

The MS was operated in positive ion mode, using Agilent MassHunter software in a neutral loss scan (NLS) setting. At collision energies of 10 eV and 20 eV, the neutral losses of 132 (for ribose) and 146 (for methyl-ribose) were monitored within a mass range of 200-500 Da. The NLS analyses used 5 µg hydrolysed tRNA.

### LC-MS data processing and analysis

As illustrated in **Scheme 1**, the UV and LC-MS/MS raw data is batch processed (n=96, by plate) using Masshunter Qualitative Analysis (Agilent, Version 8.0) and Masshunter Quantitative Analysis (Agilent, Quant-My-Way version 10.1) separately, and transformed to .csv files. If not otherwise specified, Mass data possessing method employs following parameter: Agile2 integrator algorithm, peak filter of 300 counts, left/right RT Delta 1 min, noise algorithm peak-to-peak, noise SD multiplier of 5 min, S/N 5, Accuracy Max 20% max %Dev, and smoothing function is off. For peaks with distortions (cmo^5^U, ho^5^U) and for co-eluting isobaric isomers (m^4^C/m^5^C, m^1^G/m^2^G, I/A, ho^5^U/s^2^C) the left/right RT Delta is reduced according to the RT difference. The output data from Masshunter underwent an immediate review using R to detect any shifts in retention time (RT). The script identified instances where the RT exceeded the mean value for the plate by more than 0.2 min. These cases were then flagged for a manual assessment in Masshunter to verify the precision of the peak selection process. This step is crucial for ensuring the reliability of the data before proceeding with further analysis.

An R script developed for data processing iterates through each MS data file in the directory, performing a series of steps on the data: (1) Data cleanup and extraction: For each file, it extracts MS and UV signals, and identifies samples with UV variation > 80% of the mean of each 96-sample. These samples were subjected to manual review and subsequently excluded from further analysis to prevent bias in LC-MS results due to excessively divergent sample input. (2) Normalization and fold-change calculation: The raw MS peak area of each modification (rM) is normalized by the sum of canonical ribonucleosides (rN) UV signals 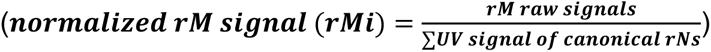, in each experiment as a control for equal sample loading into the instrument. To be noted, the normalized MS data do not reflect absolute abundance and cannot be compared directly between experiments run at different times. Then, a list of modifications existing in WT strain were subsequently processed to calculate fold changes. To compensate the signal drift over the analysis time course of each 96-well plate, the fold change was calculated in a row-based manner that was analyzed in 2 hours using following equation: 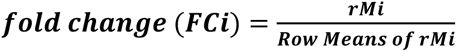. To minimize the impact of extreme values, any values more than 200% or less than 50% of the initial row means were omitted before recalculating the final row means, which then form a baseline for fold change computation for all samples in this row. (3) Annotation and data output: The normalized peakarea and fold change results were cross-referenced with mutant details (such as Gene Name, Gene ID, etc.) and saved as CSV files for further analysis. (4) Data filtering and evaluation: Mutants displaying substantial fold changes in any modification (> 2 or < 0.5) have been identified for subsequent network analyses. Furthermore, mutants with over 10 modifications exhibiting a fold change > 1.5 or < 0.7 were manually reviewed to preclude false positives in biological findings, potentially introduced by LC-MS technicalities that could result in an apparent overall upregulation or downregulation of RNA modifications.

### Arbitrary PCR and DNA sequencing

Specific PA14 mutants from frozen glycerol stocks was cultured on LB agar plates using quadrant streaking method. These plates were incubated at 37 °C overnight. For each mutant, two single colonies were picked into 3 mL LB medium (either 15 µg/mL gentamycin or 200 µg/mL kanamycin) in 15 mL falcon tube and grown at 37°C overnight (300 rpm). The overnight culture was transferred into a new 1.5 mL tube and stored at -20 °C for 1 h. The tubes were thawed and incubated at 99 °C for 10 min to lyse the cells. The cell lysate was homogenized by pipet up and down and spun at 3500 rpm for 5 min to pellet cell debris. Two-step arbitrary PCR was performed by following the instructions in the website for the “PA14 Transposon Insertion Mutant Library”^18^ (https://pa14.mgh.harvard.edu/cgi-bin/pa14/home.cgi). The PCR products were sent for Sanger sequencing (Axil Scientific, Singapore), using PMFLGM.GB-4a (5’-GACCGAGATAGGGTTGAGTG-3’) as the sequencing primer.

### Bioinformatics

Database and tools including Modomics^82^, Uniprot^104^, BV-BRC^105^, NCBI^106^, HHpred^107^ and PA14 mutant library^18^ (https://pa14.mgh.harvard.edu/cgi-bin/pa14/home.cgi) were routinely used for bioinformatic analyses. Modification and protein network were produced in Cytoscape v3.10^108^. Heatmap and clustering were performed using package “ComplexHeatmap”^109^ in R. Protein-protein interaction network was produced using STRING database^55^, with setting “interaction sources: experiments, co-expression, neighborhood, gene fusion and co-occurrence; minimal confidence score: 0.5; interactors: query protein only.” Protein IDs used in this study are listed in Supplementary **Table S8**.

### Multiple sequence alignment and structure alignment

Sequence alignment was performed using MUSCLE^110^ and viewed in Jalview^68^. Amino acids were colored according to physicochemical properties. The structure of PA14_14340, PA14 RlmN, PA14_68100, PA14_40730, PA14_16930, PA14_17650 and TapT_Ec_ were generated by SWISS-MODEL^111^ and Alphafold^112^. The protein structure models were then subjected to NPDock (Nucleic acid-Protein Dock)^113^ using the default setting with the structure of *E.coli* tRNA^Asp^ (PDB: 6UGG), tRNA^Glu^ (PDB: 5HR6), and tRNA^Val^ (PDB: 7EQJ) for the prediction of the protein-RNA complex structure. Briefly, a low-resolution method GRAMM was used to generate 20,000 alternative models (decoys) with physically reasonable geometric compatibility between protein and RNA structures. Then, the decoys were scored and clustered according to their mutual similarity, to retain groups of very similar decoys. The overall best-scored complex, as well as a representative of the largest cluster of well-scored decoys, was selected to present. Structure was analyzed and visualized using PyMol (version 2.5).

### Phylogenetic analysis

For the tree of representative bacteria, a maximum likelihood tree of 10 concatenated ribosomal proteins was generated as previously described^114^. For the tree of RlmN and PA14_40730, sequences of 5,965 RlmN and 136 PA14_40730 proteins were obtained by BLASTp search (cutoff: percentage of identity 30%; E-value 1e^-20^) in 6,616 representative bacterial genomes in BV-BRC database as collected in Jan 2021. METTL3 proteins in human, mouse and drosophila were used as the out group. For the tree of PA14_68100, sequences of 1460 TrmH and 340 PA14_68100 proteins were obtained by BLASTp search. The obtained protein sequences were aligned using MUSCLE^110^ and trimmed by BMGE v1.12^115^. The tree is inferred using FastTree^116^ with default parameters and 100 bootstrap replicates generated with SeqBoot^117^. The trees are visualized using iTOL^118^. Branches are colored by phylogenetic affiliation at phylum level.

## Data availability

The mass spectrometry data have been deposited to the ProteomeXchange Consortium via the PRIDE partner repository with the dataset identifier PXD053297 (http://www.ebi.ac.uk/pride). Sequencing data have been deposited in the NCBI SRA database with BioProject ID PRJNA1126677.

## Code availability

All data analysis and data visualization R scripts are available in the Github repository at https://github.com/jingjsunny/tRNAmodi.git.

## Acknowledgments

We thank the MIT BioMicro Center and its Director, Dr. Stuart Levine, for support and advice during the performance of the studies presented here. The authors gratefully acknowledge funding from the Singapore National Science Foundation under the Singapore-MIT Alliance for Research and Technology Antimicrobial Resistance Interdisciplinary Research Group (PCD), NIH Transformative Award ES031576 (PCD), and Center grant P30-ES002109 from the National Institute of Environmental Health Sciences of the National Institutes of Health.

