## Supplementary Information for "tRNA modification profiling reveals epitranscriptome regulatory networks in *Pseudomonas aeruginosa*"

### *Contents*


- **Scheme 1.** Workflow for LCMS data process.
- **Supplementary Figure 1**. Validation of 4 M GITC cell lysis protocol for Pseudomonas aeruginosa.
- **Supplementary Figure 2.** High-throughput tRNA modification profiling platform application on human embryotic kidney cells and mouse brain.
- **Supplementary Figure 3.** PEG-8000 and salts effect on beads size exclusion in RNA species in PA14.
- **Supplementary Figure 4.** Scheme of magnetic beads-based RNA purification workflow and the deck layout of Tecan EVO150.
- **Supplementary Figure 5.** Reproducibility of RNA purity check of 96 tRNA samples purified by Tecan on a bioanalyzer pico chip.
- **Supplementary Figure 6.** Command optimization on the EVO 150.
- **Supplementary Figure 7:** Optimizing chromatography performance.
- **Supplementary Figure 8.** LC-MS/MS performance characteristics.
- **Supplementary Figure 9.** Optimizing RNA sample processing conditions.
- **Supplementary Figure 10.** High-resolution mass spectrometry confirmation of signals (m6t6A and ms2io6A) found by neutral loss scan in PA14 WT strain.
- **Supplementary Figure 11.** Visualization of RNA-Seq coverage across the aroB region of PA14 WT, aroB mutant PA14NR:42535 and aroB mutant PA14NR:38358.
- **Supplementary Figure 12.** Changes in levels of the i6A family of position 37 tRNA modifications dominate the PA14 knockout library.
- **Supplementary Figure 13.** Functional annotation of genes for RNA-modifying enzymes.
- **Supplementary Figure 14.** Sequence alignment of PA14_14340 and RlmF protein sequences.
- **Supplementary Figure 15.** Maximum-likelihood phylogenetic tree of 5965 RlmN and 136 PA14_40730 like proteins.
- **Supplementary Figure 16.** Sequence alignment of selected RlmN-like, and 40730-like protein sequences.

- **Supplementary Figure 17.** Sequence alignment of PA14_68100 homologs.
- **Supplementary Figure 18.** Taxonomic and phylogenetic analysis of TrmH and PA14_68100-like proteins.
- **Supplementary Figure 19.** Annotation of PA14_16930 as CsdA involved in ct6A formation.
- **Supplementary Figure 20.** The alignment of PA14_17650 and $TapT_{Ec}$.
- **Supplementary Table 1:** Configuration of Tecan EVO150 parameters used for magnetic beads-based tRNA isolation from crude lysates.
- **Supplementary Table 2:** *Separate spreadsheet.*
- **Supplementary Table 3:** Detect of limit (LOD) and Quantification of limit (LOQ) of modified ribonucleosides using the rapid UHPLC/MS method developed in this study. *Separate spreadsheet.*
- **Supplementary Table 4:** Modification table*. Separate Excel spreadsheet.*
- **Supplementary Table 5:** Screening results of whole library. *Separate spreadsheet.*
- **Supplementary Table 6.** Verification of known tRNA modifying enzymes. *Separate spreadsheet.*
- **Supplementary Table 7.** Gene pathway list. *Separate spreadsheet.*
- **Supplementary Table 8.** Protein IDs used in this study*. Separate spreadsheet.*

*Supplementary Figures*

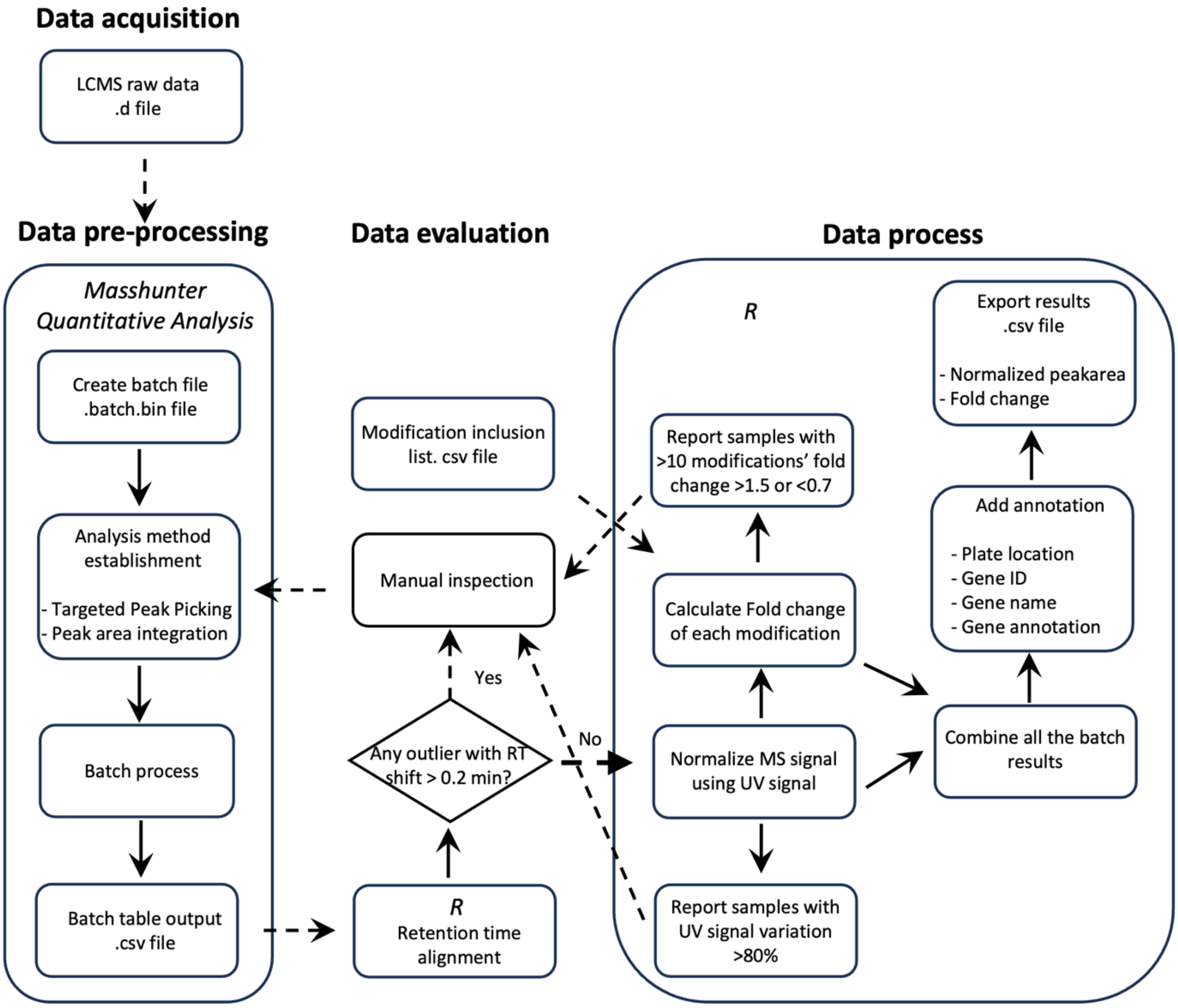


**Supplementary Scheme 1. Workflow for LCMS data process.** It consists of three main sections: (1) Batch processing of raw UV and MS data using the Masshunter software (Agilent, USA). This step involves target peak integration, identification. (2) Data evaluation and curation using R. (3) Normalization of MS data based on UV signals of canonical ribonucleosides, including followed by statistical analysis.

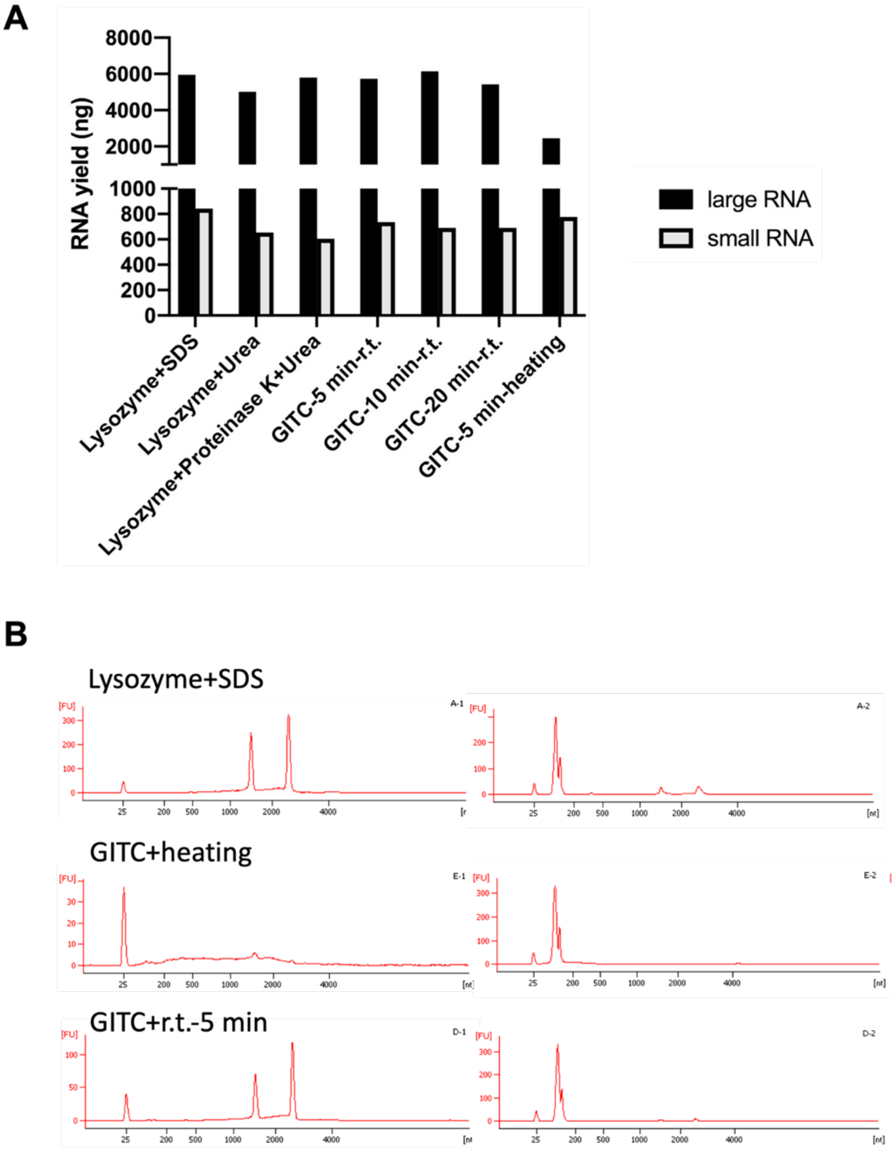


**Supplementary Figure 1. Validation of 4 M GITC cell lysis protocol for *Pseudomonas aeruginosa*.** A. RNA yield assessment yielded from 0.3 $OD_{600}$ cells using different cell lysis protocol. Large RNA and small RNA are separated using magnetic beads-based method described in this study. From left to right: Lysozyme (10 mg/mL), SDS (5%) in TE buffer, pH 8.0, room temperature, incubate 5 min (commercial RNA extraction kit protocol for bacteria); Lysozyme (10 mg/mL), Urea (2 M) in TE buffer, pH 8.0, room temperature, incubate 5 min; Lysozyme (10 mg/mL), Proteinase K (0.1 mg/mL), Urea (2 M) in TE buffer, pH 8.0, room temperature, incubate 5 min; GITC (4 M) in Tris buffer, pH 8.0, room temperature, vertexing 5 min/10 min/20 min (1500 rpm); GITC (4 M) in Tris buffer, pH 8.0, 65 ℃, incubate 5 min. B. RNA integrity assessment on Agilent bioanalyzer pico chip (25-4000 nt).

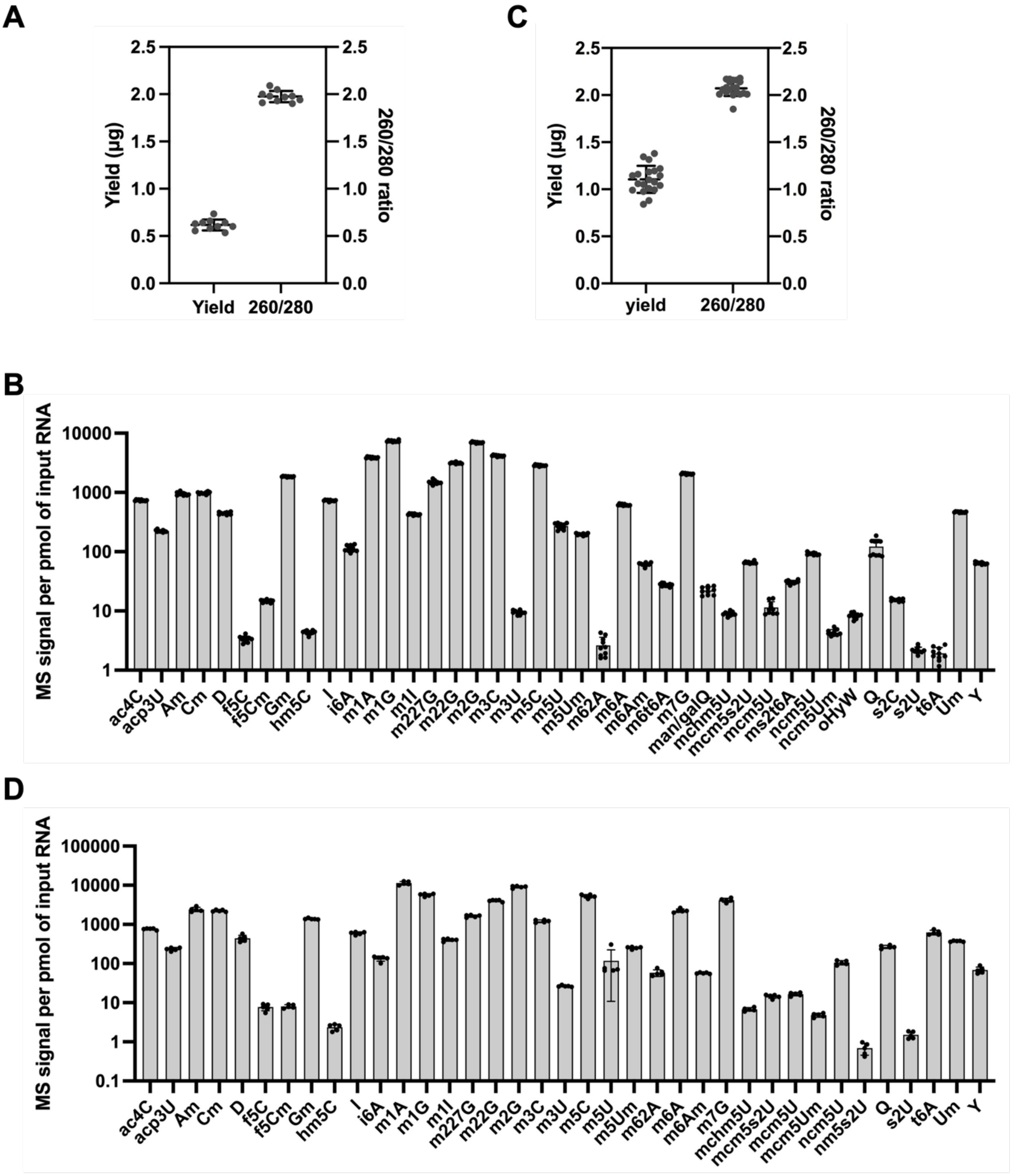


**Supplementary Figure 2. High-throughput tRNA modification profiling platform application on human embryotic kidney cells and mouse brain.** Data are plotted with 2 biological replicates (n=10) from (A and B) GripTite 293 MSR and (C and D) mouse C57BL/6 brain (9-month-old, n=5 mice). The level of each modification was presented as $\log_{10}$ scale of MS signal per pmol of input hydrolyzed RNA.

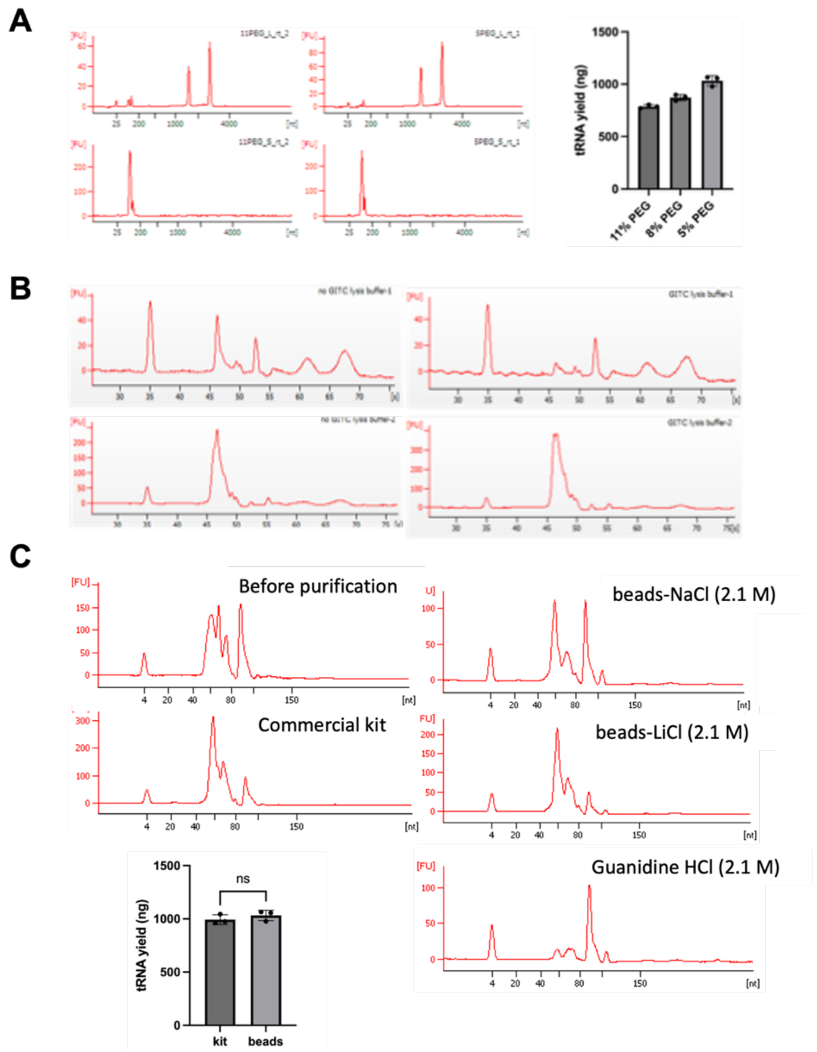


**Supplementary Figure 3. PEG-8000 and salts effect on beads size exclusion in RNA species in PA14**. (A) Bioanalyzer pico chip traces of isolated RNA species in presence of 11% PEG-8000 (Left upper: 1st round beads elutes; Left lower: 2nd round beads elutes) and 5% PEG-8000 (Middle upper: 1st round beads elutes; Middle lower: 2nd round beads elutes). Right: tRNA yield in presence of 11% vs. 5% PEG-8000. (B) Bioanalyzer small chip traces of large RNA species and small RNA species purified by beads in absence (Left upper: elutes from 1st round beads; Left lower: elutes from 2nd round beads) and presence of GITC (Right upper: elutes from 1st round beads; Right lower: elutes from 2nd round beads). The results indicated that the presence of GITC helps improve the resolution in separation of 5s rRNA and tRNA. (C) Bioanalyzer small chip traces of total RNA before purification (left upper), small RNA species purified by commercial spin column kit (left lower), beads in 2.1 M NaCl (right upper), 2.1 M LiCl (right middle) and 2.1 M GuCl (Right lower). The effect of pH is also explored (data not shown). Under acidic condition (pH 5.3), carboxyl-magnetic beads lose the ability to bind RNA, while basic condition (pH 8.0) slowly causes RNA hydrolysis. A close to neutral condition (pH 7.5) is chosen for all the assays in this study. The tRNA yield from magnetic beads-based method is comparable with commercial kit. *P* value is determined by t-test, n=3.

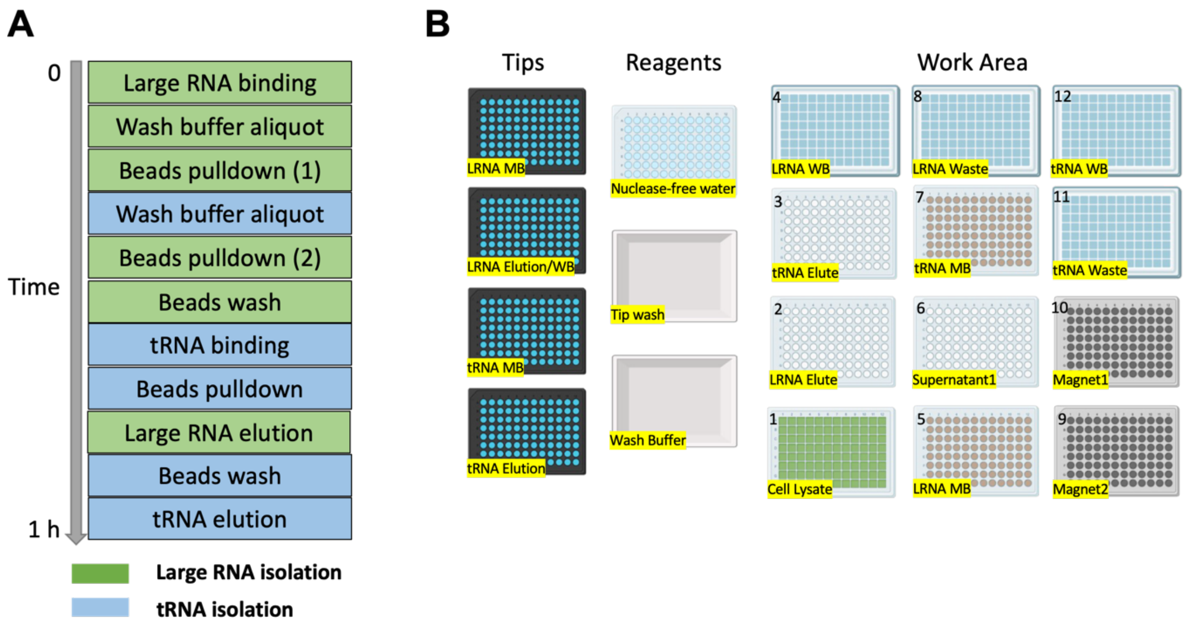


**Supplementary Figure 4. Scheme of magnetic beads-based RNA purification workflow and the deck layout of Tecan EVO150.** (A) Scheme of magnetic beads-based RNA purification workflow by Tecan EVO150, involving both large RNA and tRNA purification. The tasks are performed in the presented sequence, and the entire procedure is completed within one hour. (B) The deck layout of Tecan EVO150. Different icons represent different types of labware used throughout the RNA purification process. The specific application of each labware is highlighted. The deck position is indicated by the number labeled on the left top of labware.

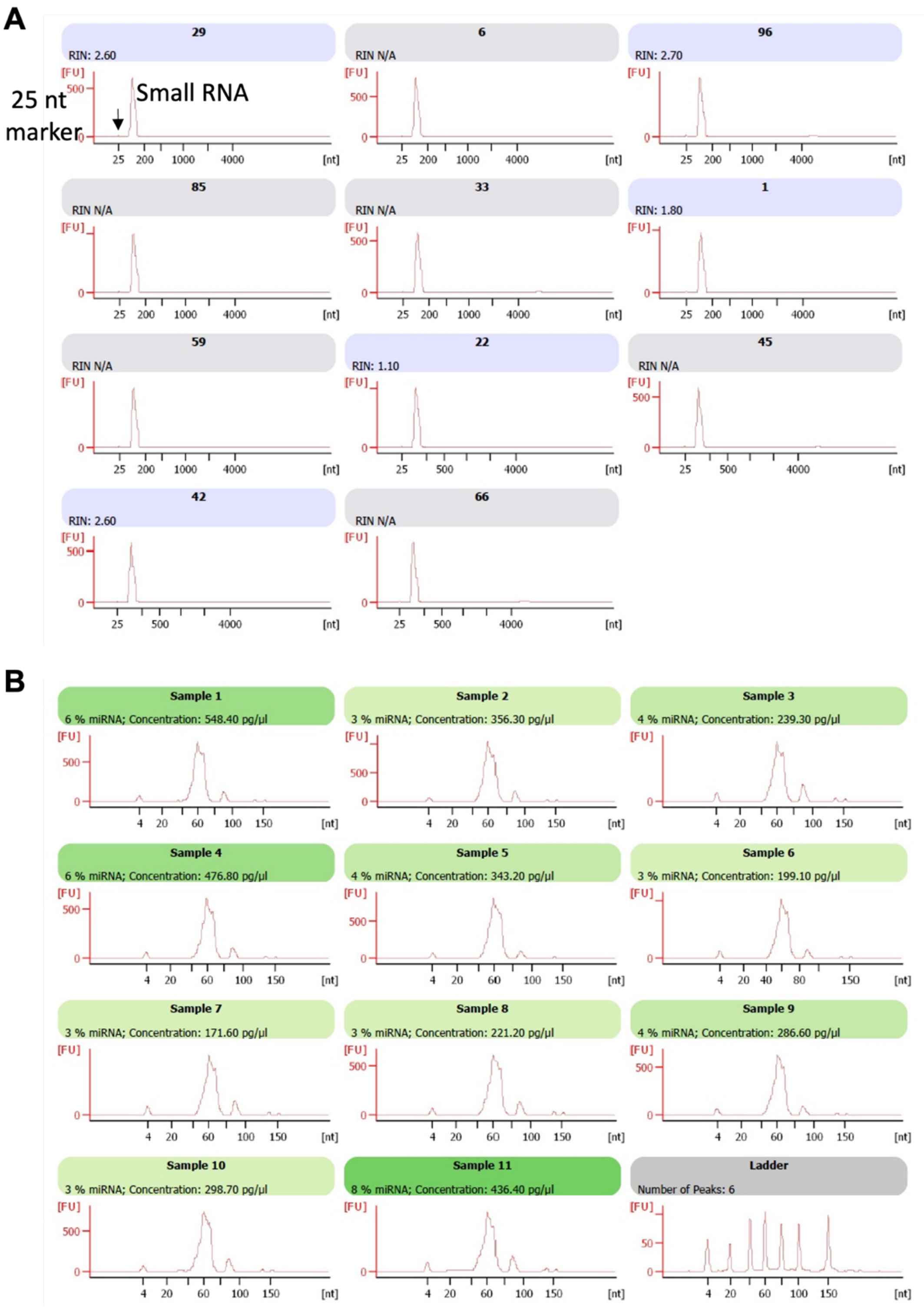


**Supplementary Figure 5. Reproducibility of RNA purity check of 96 tRNA samples purified by Tecan on a bioanalyzer pico chip.** (A) and small chip (B). The RNAs were overloaded to check potential rRNA contamination. All of yielded tRNAs are free of large rRNA and with consistent, small traces of 5s rRNA, which is unlikely resulting bias in modification analysis.

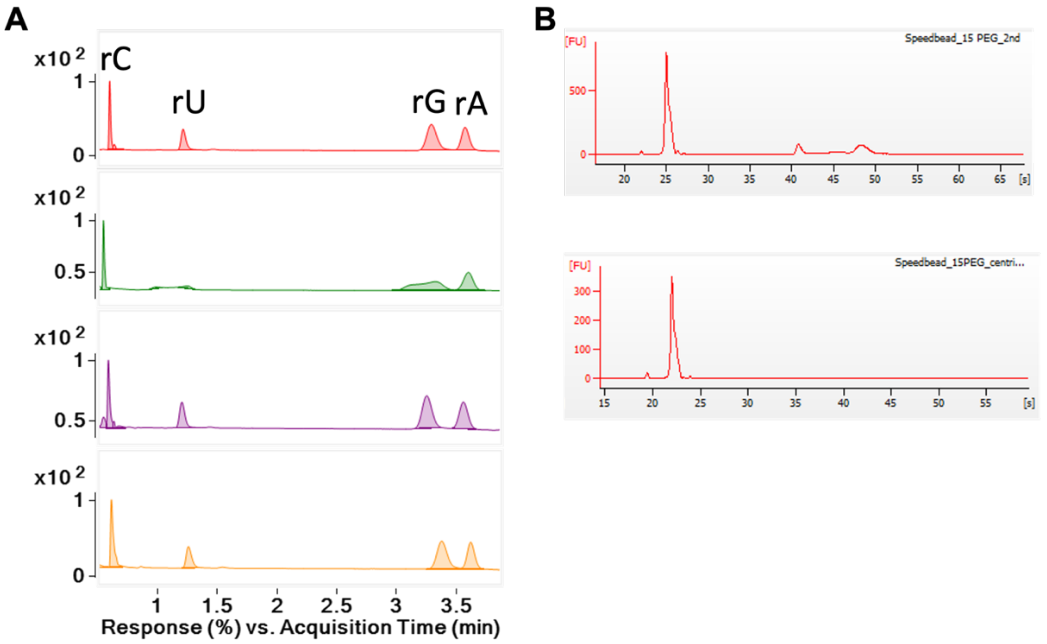


**Supplementary Figure 6. Command optimization on the EVO 150** to (A) minimize peaks from incomplete buffer removal during tRNA isolation. Optimization on command of EVO 150 to avoid peak shape deteriorated during LCMS analysis derived from beads wash buffer leftover derived from tRNA isolation. From top to bottom: UV chromatograph of hydrolyzed tRNA purified manually, automatically by EVO 150 before command optimization, dried sample B by speedvac and after command optimization; (B) eliminate rRNA contamination induced by 1$^{st}$ round beads carryover. From top to bottom is the bioanalyzer traces on a pico chip: purified tRNA yielded from 1$^{st}$ round beads pulldown by magnet twice; beads pulldown by magnet twice and with a plate centrifugation in between.

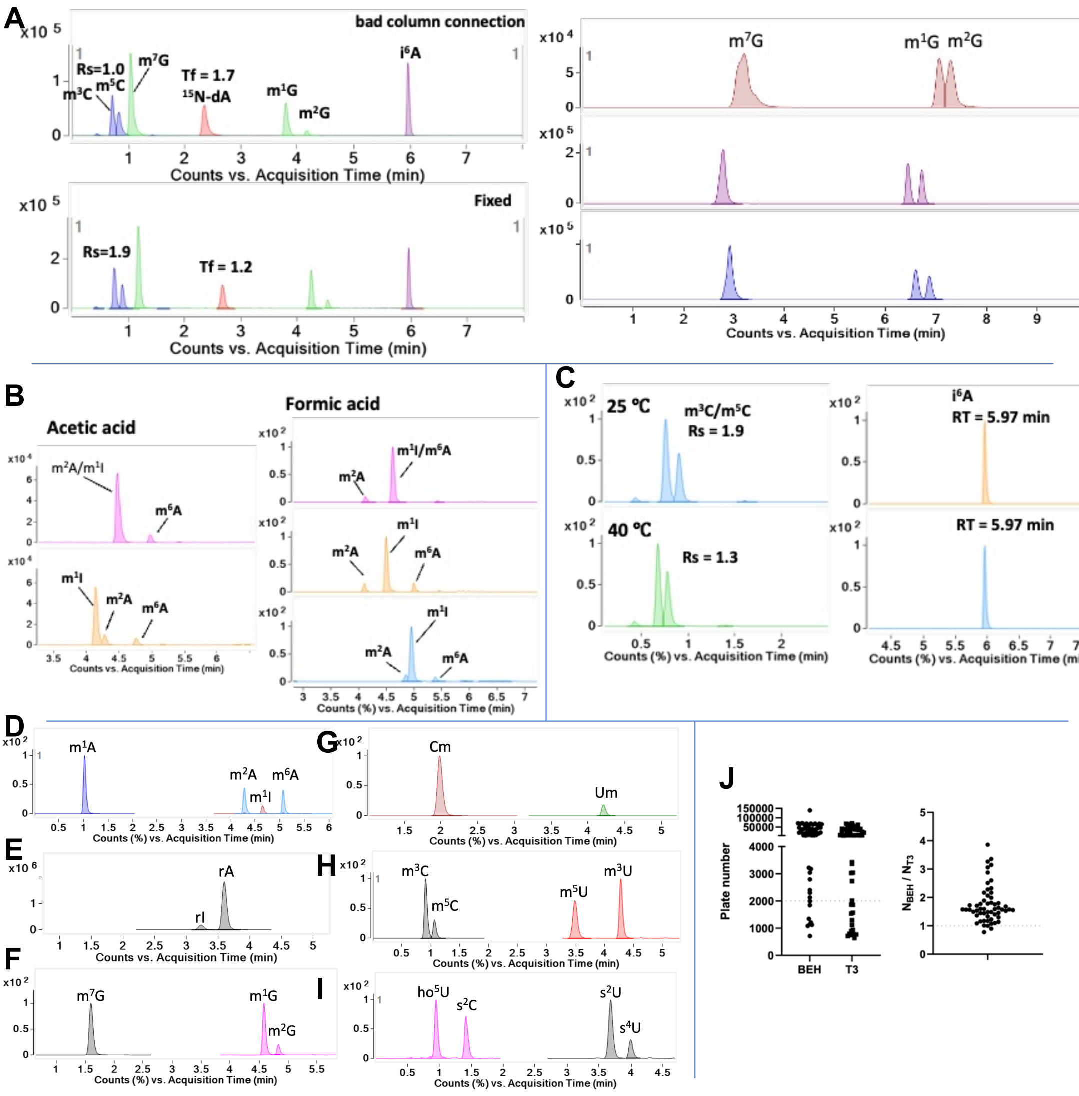


**Supplementary Figure 7. Optimizing chromatography performance.** (**A**) Minimizing column dead volume to improve peak resolution. *Left*: Dead volume due to longer tubing and larger connection induces peak tailing and reduces peak resolution (upper), solved in left/lower; *Right:* Dead volume in standard 10 mm flow cell (13 µL; upper) reduces peak resolution, solved with a lower volume bypass UV detector (middle) or high pressure 6 mm flow cell (1.7 µL). (**B**) Optimizing mobile phase pH to resolve m$^1$I, m$^2$A and m$^6$A by dynamic MRM due to similar transitions (m$^1$I *m/z* 283>151, m$^2$A/m$^6$A *m/z* 282>150). *Acetic acid:* top, 0.1%; bottom, 0.02%. *Formic acid:* top, 0. 1%; middle, 0.02%; bottom, 0.01%. 0.02% FA offers best separation. (**C**) Optimizing column temperature for m$^3$C/m$^5$C (left) and i$^6$A (right). *Upper*: 25 ºC. *Lower*, 40 ºC. 25 °C offers separation of isobaric m$^3$C and m$^5$C without slowing elution. (**D-I**) Chromatography of ribonucleosides with the same or similar CID transitions. (**D**) m$^1$A, m$^2$A, m$^6$A: *m/z* 282>150, m$^1$I *m/z* 283>151. (**E**) rI, *m/z* 269>137; rA, *m/z* 268>136. (**F**) m$^7$G, m$^1$G and m$^2$G: *m/z* 298>166; (**G**) Cm, *m/z* 258>112; Um, *m/z* 259>113; (**H**) m$^3$C, m$^5$C: *m/z* 258>126; m$^3$U, m$^5$U: *m/z* 259>127; (**I**) ho$^5$U, *m/z* 261>129; s$^2$C, *m/z* 260>128; s$^2$U, s$^4$U: *m/z* 261>129. (**J**) The Waters BEH C18 column (50 mm, 2.1 mm, 1.7 µm) was chosen over the Waters HSS T3 (100 mm, 1 mm, 1.8 µm) column for shorter run times and better resolution, reflected in the larger number of theoretical plates (plate number, N). BEH conditions: 0.35 mL/ min, 330 bar, 25 °C, 100% solution A (water, 0.02% formic acid) for 2 min, followed by 2-4 min at 0%-8% solution B (70% acetonitrile, 0.02% formic acid), and from 4-5.9 min at 8%-100% B. TSS conditions: 0.25 mL/ min, 650 bar, 40 °C, 0-1.5 min, 0-3% solution B (70% acetonitrile, 0.02% formic acid); 1.5-5 min, 3-14% B; 5-5.10, 14-100% B; 5.1-8 min, 100 % B.

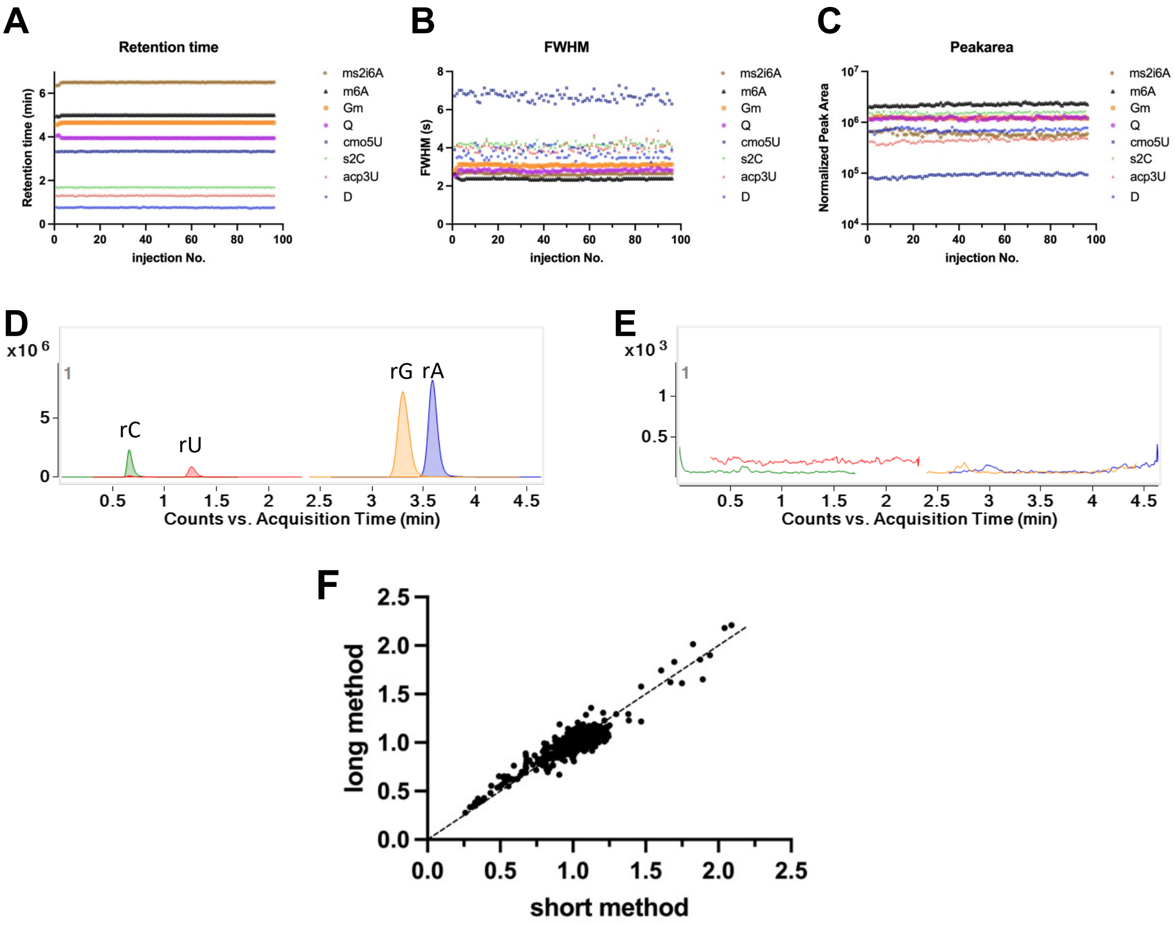


**Supplementary Figure 8. LC-MS/MS performance characteristics.** (**A-E**) LCMS method intra-day assessment using the signals of RNA modifications in a 200 ng hydrolysed PA14 tRNA injection (n=96). Eight modifications are shown as representatives in this figure due to their retention times spanning the entire analyte elution window. The statistical data in Results and Discussion is calculated based on all modifications detected in the PA14 WT strain. The method performance of 96 injections is assessed by the signal stability of retention time (**A**), full width at half maximum (FWHM, **B**) and normalized peak area (**C**). (**D,E**) Between sample injections, blank injections were performed to assess signal carryover. (**D**) Representative sample injection and (**E**) subsequent blank injection following the sample. (**F**) Correlation comparison in results derived from same sample matrix using short LCMS method vs. long LCMS method. Both axis represent the level fold change of modifications detected in 293T cell samples (n=16). The LC condition for long method: a Phenomenex Synergi Fusion-RP C18 column (100 × 2.0 mm, 2.5 µm) is coupled to an Agilent 1290 HPLC system at 35 °C and a flow rate of 0.35 mL/min, with a gradient starting with 100% solution A (5 mM ammonium acetate, pH 5.3), followed by 0-10% solution B (acetonitrile) 0-10 min; 10%-40% solution B, 10-14 min; 40%-80% solution B, 14-15 min; 80%-90% solution B, 15-15.1 min;  90% solution B, 15.1-18 min.

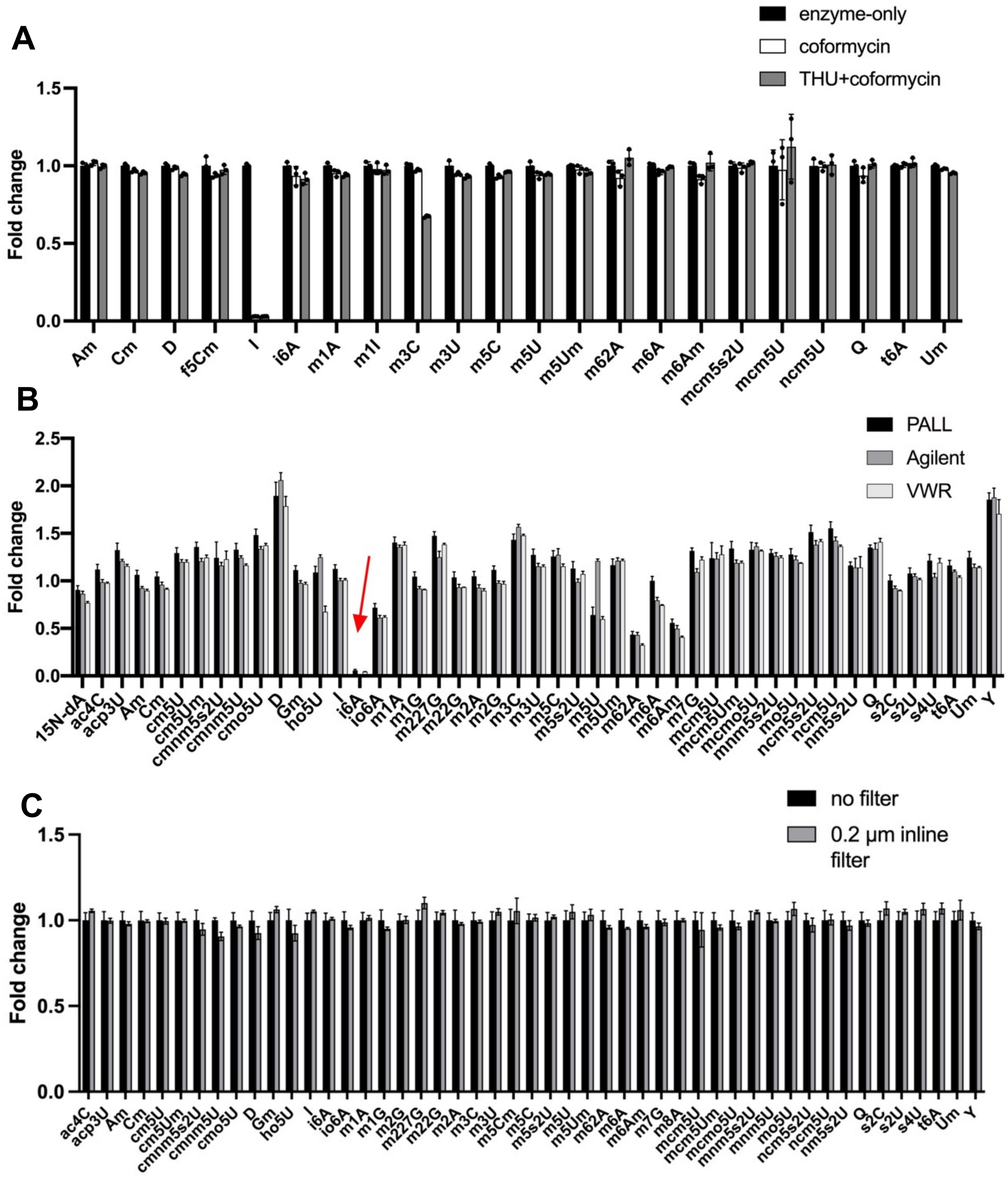


**Supplementary Figure 9. Optimizing RNA sample processing conditions.** (**A**) The effect of additives in tRNA hydrolysis enzyme cocktails on RNA modification analysis. tRNA extracted from HEK 293 cells (5 µg) was hydrolyzed in 3 different enzyme master mixes: _Enzyme-only_: 5 mM Tris, 2.5 mM $MgCl_2$, 12.5 U benzonase, 5 U CIAP, and 0.15 U PDE I; _Coformycin:_ 5 mM Tris, 2.5 mM $MgCl_2$, 12.5 U benzonase, 5 U CIAP, 0.15 U PDE I, and 5 ng coformycin; _THU+coformycin_: 5 mM Tris, 2.5 mM $MgCl_2$, 12.5 U benzonase, 5 U CIAP, 0.15 U PDE I, 5 ng coformycin, and 50 ng tetrahydrouridine (THU). (**B,C**) The impact of filters for ribonucleosides LCMS analysis. (**B**) Fold-change values for LC-MS/MS analysis of synthetic standards after filtration of tRN**B**A hydrolysates with 3 commercial 10,000 Da filters: VWR (tube format), PALL (96-well plate format), and Agilent (96-well plate format). Fold-change was calculated relative to unfiltered signals. (**C**) Fold-change values for LC-MS/MS analysis of synthetic standards processed using a 0.2 µm inline filter. No significant column clogging was observed after ~1000 injections, though enzymes cannot be removed using 0.2 µm inline filter.

**A**

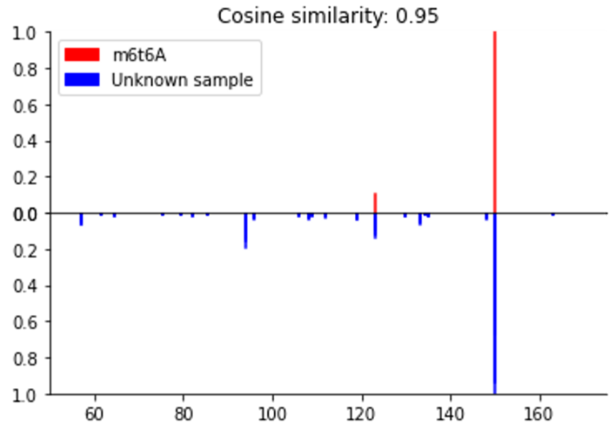


[M+H]+: 427.1577, found 427.1571, Δppm: -1.4

**B**

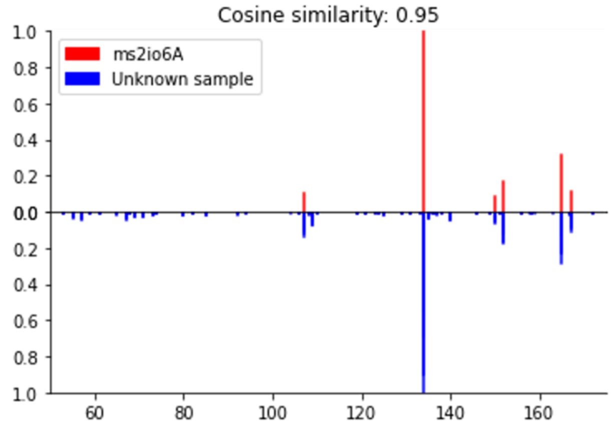


[M+H]+: 398.1498, found 398.1494, Δppm: -1.0

**Supplementary Figure 10. High-resolution mass spectrometry confirmation of signals ($m^6t^6A$ and $ms^2io^6A$) found by neutral loss scan in PA14 WT strain.** The signals of $m^6t^6A$ (left) and $ms^2io^6A$ (right) were confirmed by higher-energy collisional dissociation (HCD)-MS and spectral library matching by injecting 2 μg hydrolyzed tRNA. The LC and MS parameters strictly follow the protocol as previously reported (doi: 10.1021/acs.analchem.2c03172). No U modifications were discovered using this method.

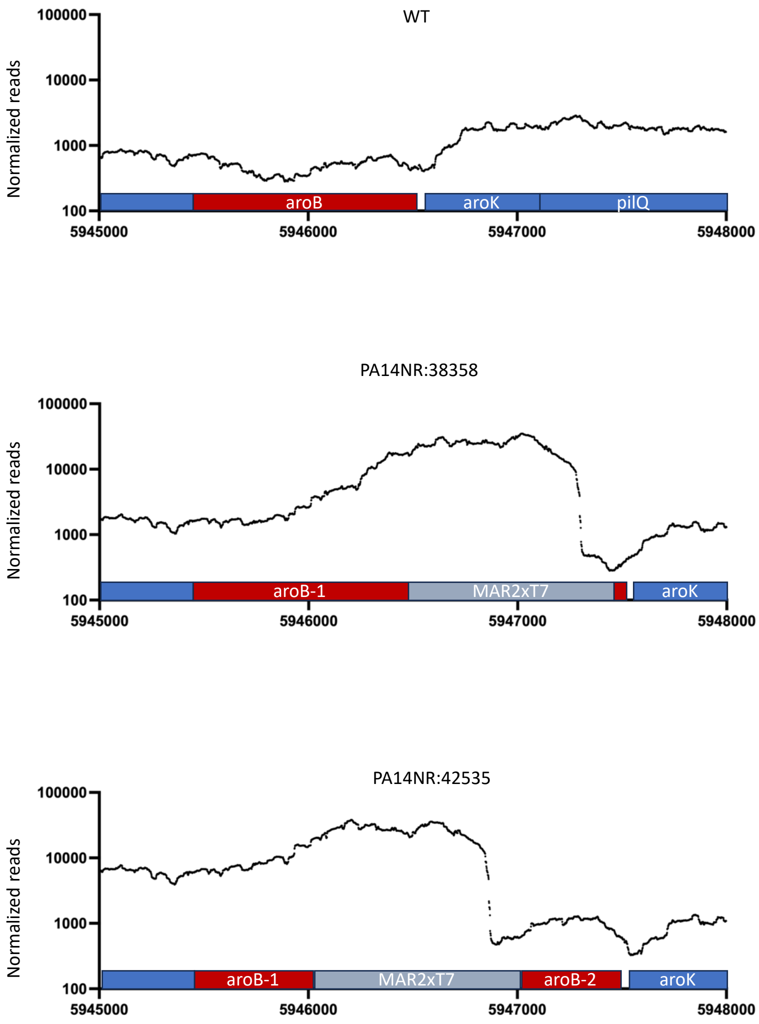


**Supplementary Figure 11. Visualization of RNA-Seq coverage across the aroB region of PA14 WT, aroB mutant PA14NR:42535 and aroB mutant PA14NR:38358.** Black curves represent read coverage, and the aroB gene is highlighted in red. The introduction of the high-expression transposon MAR2xT7 elevated expression of a truncated aroB upstream of this transposon. In PA14NR:42535, the truncated aroB has a 32bp deletion at the 5'end, potentially allowing for the synthesis of a functional N-truncated AroB protein. While in PA14NR:38358, the truncated aroB is too short to produce functional AroB protein.

**A**

| | Number per $10^6$ rA |
|---|---|
| $i^6A$ | 402 ± 14 |
| $io^6A$ | 84.5 ± 3.0 |
| $ms^2i^6A$ | 910 ± 44 |
| $ms^2io^6A$ | 3482 ± 54 |

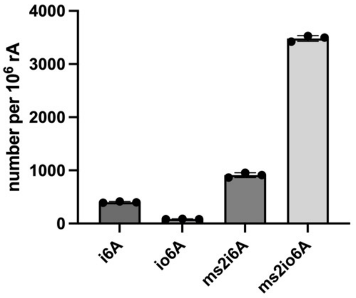


**B**

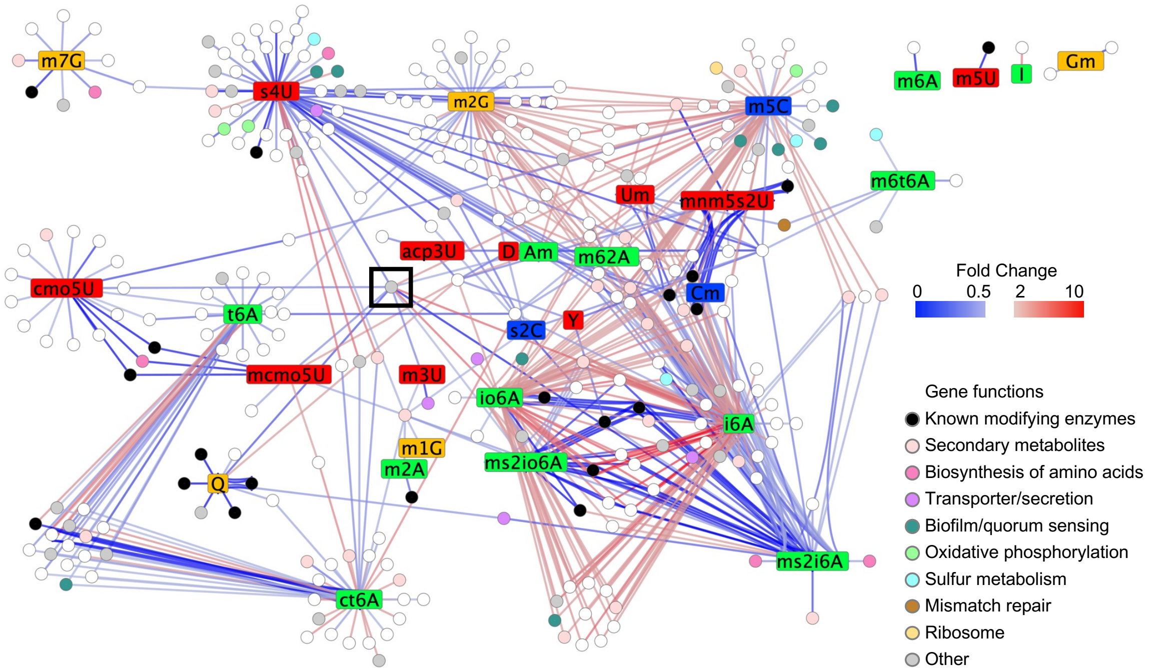


**Supplementary Figure 12. Changes in levels of the $i^6A$ family of position 37 tRNA modifications dominate the PA14 knockout library.** (**A**) Levels of $i^6A$, $io^6A$, $ms^2i^6A$ and $ms^2io^6A$ expressed as number of modifications per $10^6$ adenosines (rA) in the transposon intergenic mutant control (mean ± SD for n=3). Absolute quantification of $i^6A$, $io^6A$, and $ms^2i^6A$ was achieved using external calibration curves with synthetic standards. Quantification of $ms^2io^6A$ (no standard available) was achieved by in-line UV absorbance using the extinction coefficient of $io^6A$ given their comparable molecular structures. (**B**) Protein-modification network of modifications (center square in each cluster) linked to a PA14 mutant gene (nodes/circles) that affects the modification level. Edge/line color indicates the modification fold-change (increasing red, decreasing blue), edge/line thickness correlates with the STRING database confidence score, and node/circle color indicates the GO function of the gene. For visualization, only clusters with ≥3 proteins and edges with confidence scores ≥ 0.5 are shown.

**A**

| Modification | Gene ID | Gene Name | Known function | Large RNA | Small RNA |
|---|---|---|---|---|---|
| $C_m$ | PA14_68100* | | | 0.944±0.062 | 0.735±0.024 |
| | PA14_68110* | | | 1.018±0.098 | 0.765±0.005 |
| $G_m$ | PA14_68100* | | | 0.960±0.029 | 0.007±0.001 |
| | PA14_68110* | | | 0.971±0.040 | 0.292±0.036 |
| | PA14_65190 | rlmB | 23s rRNA (2251) | 0.055±0.063 | 1.057±0.042 |
| $m^2A$ | PA14_14830 | rlmN | 23s rRNA (2508)<br>tRNA (37) | 0.024±0.026 | 0.214±0.014 |
| | PA14_40730* | | | 0.991±0.034 | 0.841±0.027 |
| $m^6A$ | PA14_14340* | rlmF | 23s rRNA (1618) | 0.548±0.003 | 0.008±0.001 |
| | PA14_66340 | rlmJ | 23s rRNA (2030) | 0.467±0.001 | 1.047±0.041 |
| $m^6_2A$ | PA14_07730 | rsmA | 16s rRNA (1518/1519) | 0.008±0.001 | 0.086±0.026 |

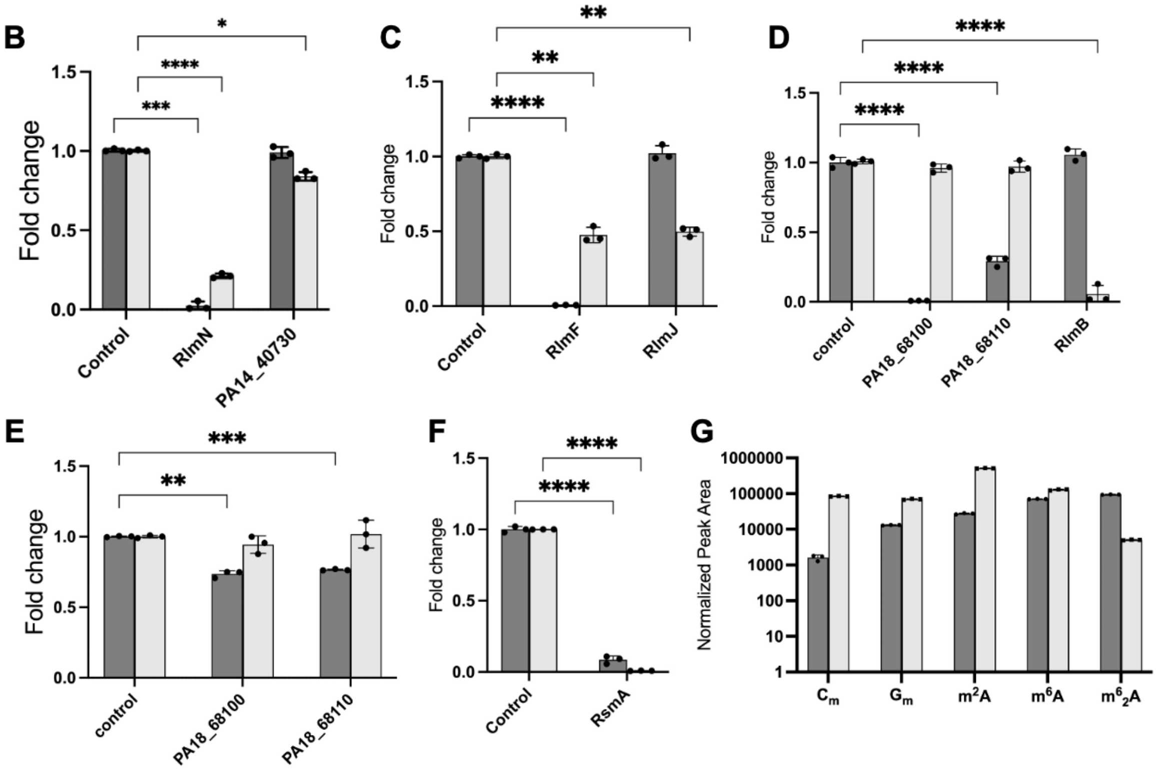


**Supplementary Figure 13. Functional annotation of genes for RNA-modifying enzymes. (A)** Summary of enzymatic functions annotated in this study in comparison to existing enzymes responsible for the same modifications. Asterisks denote newly annotated enzymes. Fold-change values for modifications were calculated relative to a transposon intergenic mutant control (n=3). **(B-G)** Analysis of modification levels in large RNA and small RNA fractions from PA14. **Dark gray**: large RNA; **light gray**: small RNA **(B)** $m^2A$ levels of PA14_14830 (RlmN) and PA14_40730 mutants. **(C)** $m^6A$ levels in RlmF and RlmJ mutants. **(D)** $G_m$ levels in PA14_68100, PA14_68110 and PA14_65190 (RlmB) mutants. **(E)** $C_m$ levels in PA14_68100, PA14_68110 and PA14_65190 (RlmB) mutants. **(F)** $m^6_2A$ levels in PA14_07730 (RsmA) mutant. Fold-change values were calculated relative a transposon intergenic mutant control. **(G)** The relative abundance of modifications in large RNA and small RNA fractions. Note the $\log_{10}$ y-axis scale. (**B-F**) Statistical significance is indicated as follows: ns (not significant), * ($p < 0.05$), *** ($p < 0.001$), and **** ($p < 0.0001$), determined by *two-way ANOVA* test.

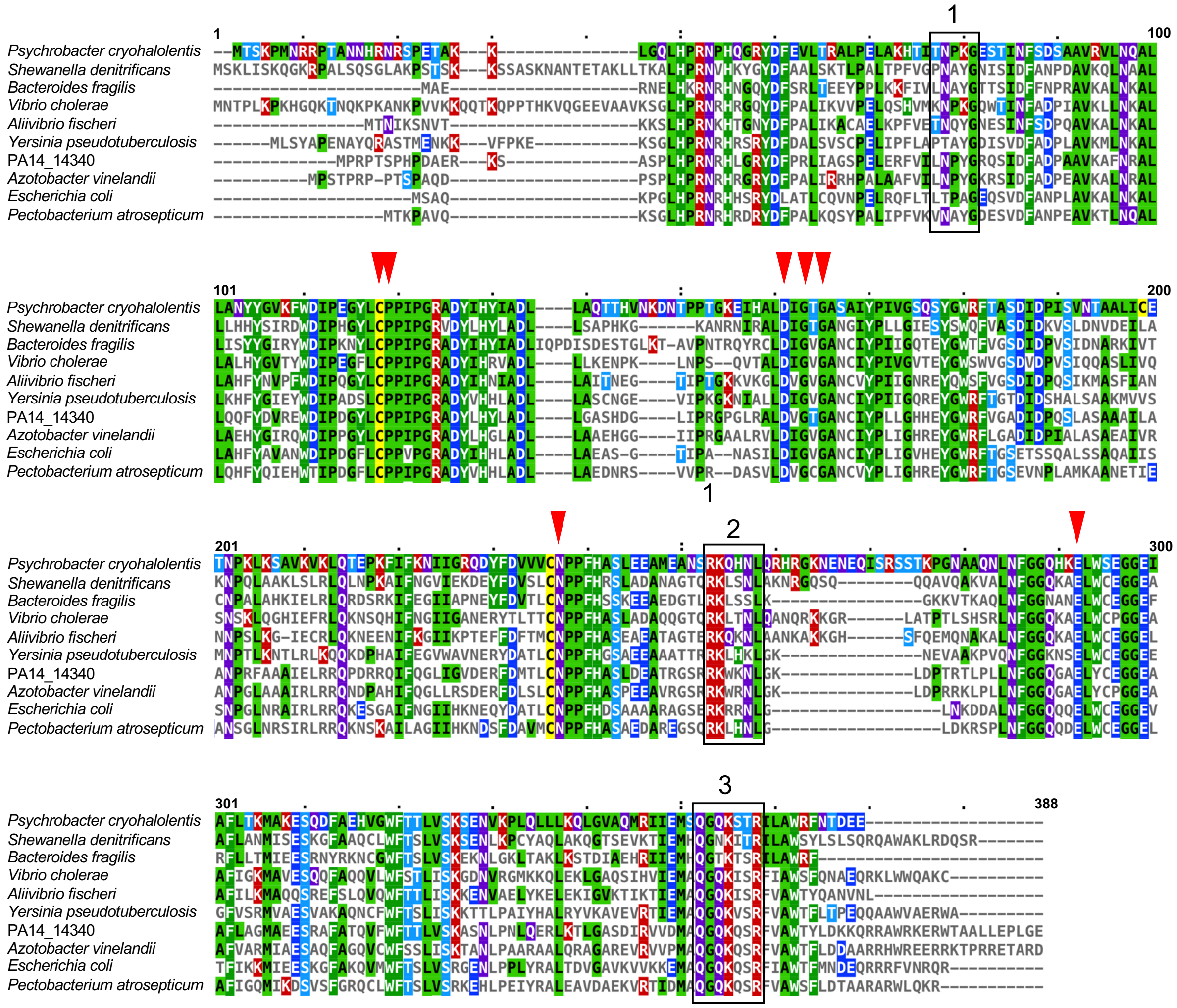

**Supplementary Figure 14. Sequence alignment of PA14_14340 and RlmF protein sequences.** Sequence alignment of PA14_14340 and 9 selected RlmF protein sequences. Amino acid coloring reflects physicochemical properties. Conserved residues in the proximity of SAH are indicated by red triangles. Three turns predicted in the PA14_14340 protein structure that fit the groove of ASL of tRNA are boxed. Dashes indicate gaps in the sequence alignment. Uniprot IDs for proteins included in multiple alignment were listed in Table S8.

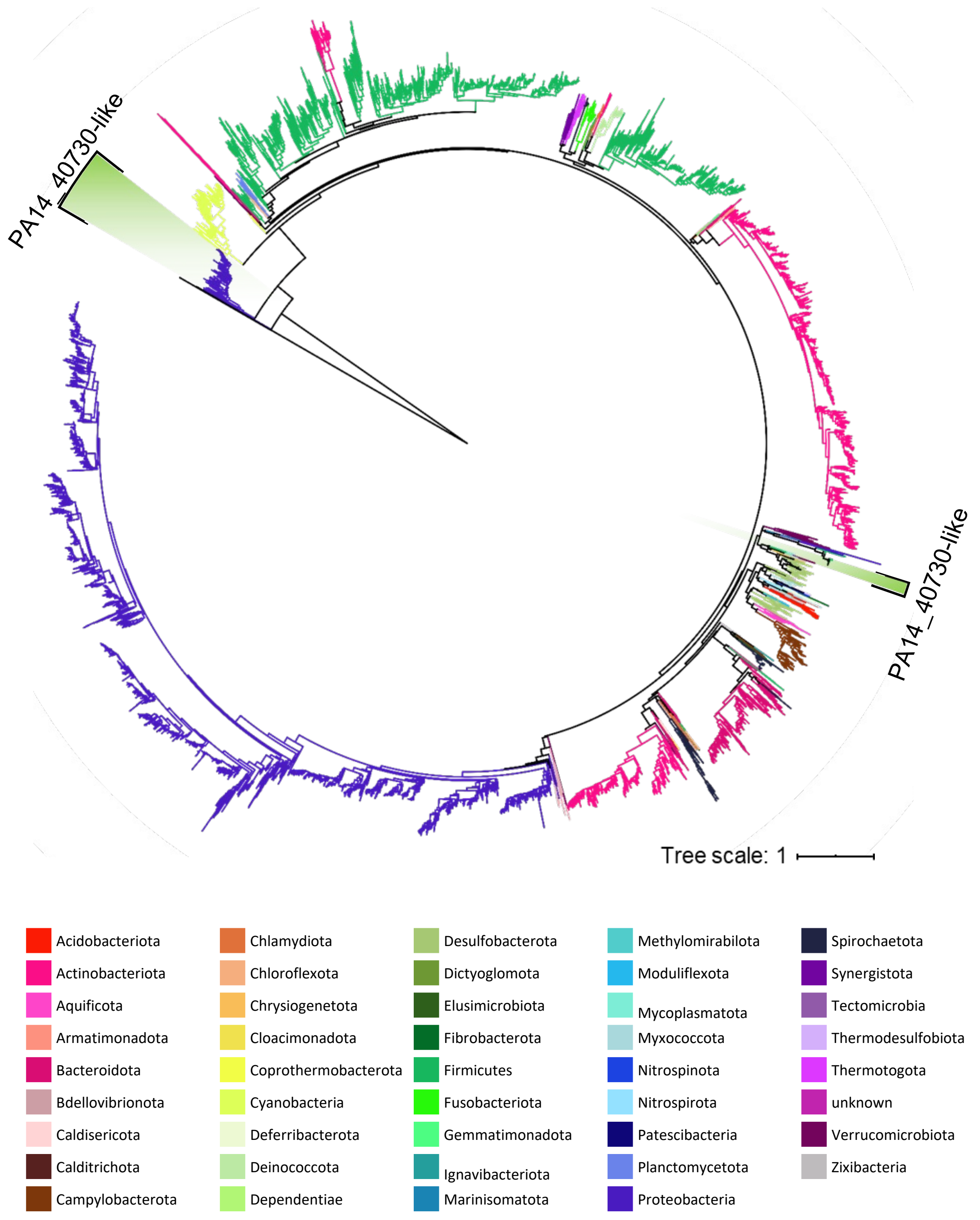


**Supplementary Figure 15. Maximum-likelihood phylogenetic tree of 5965 RlmN and 136 PA14_40730 like proteins.** Sequences are obtained by BLASTp search in 6616 representative bacterial genomes with METTL3 proteins in human, mouse and drosophila as the out group. Branches are colored by phylogenetic affiliation at phylum level. The light green sections are PA14_40730 like proteins and the rest are RlmN homologs.

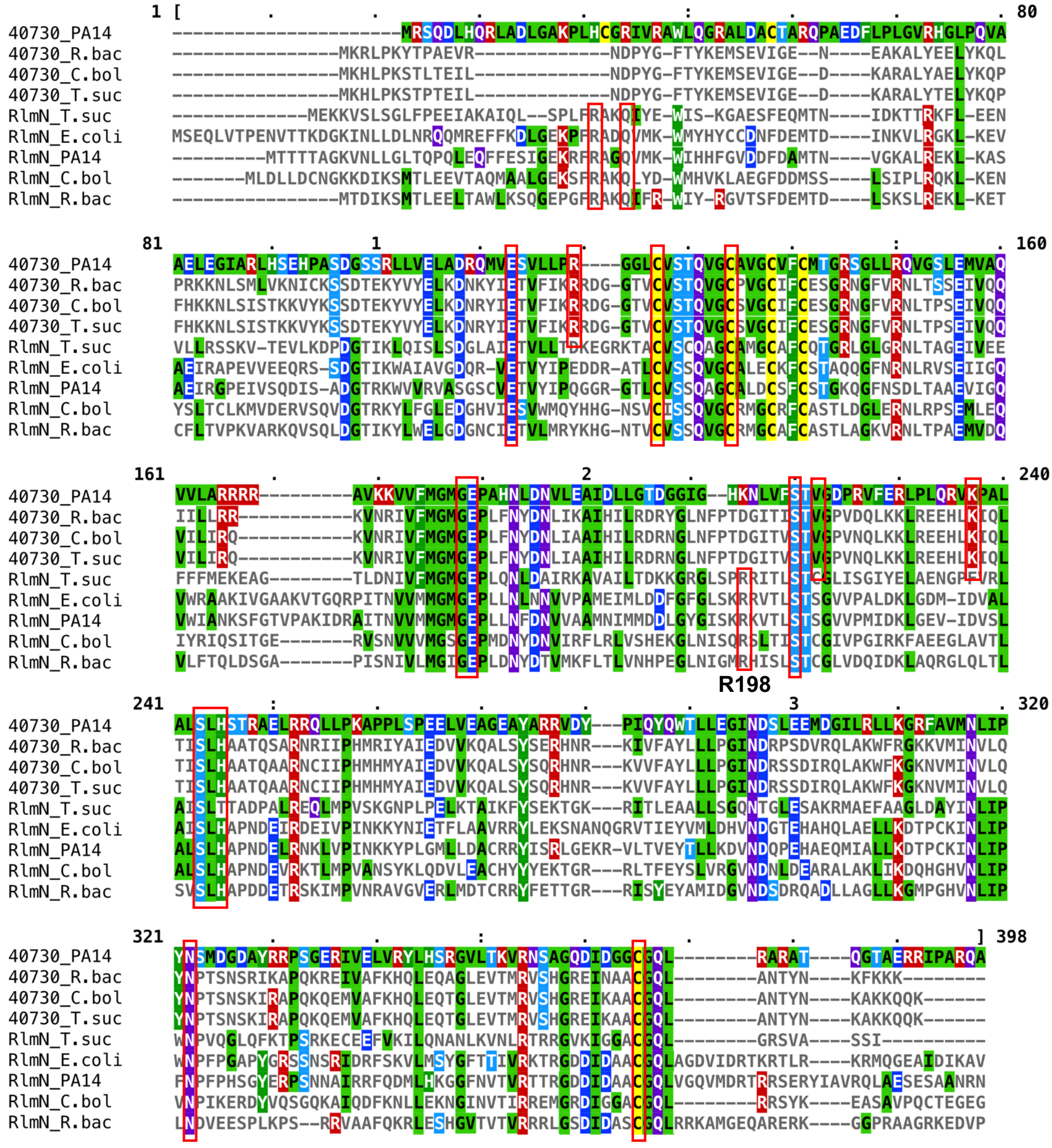


**Supplementary Figure 16. Sequence alignment of selected RlmN-like, and 40730-like protein sequences.** Sequence alignment of 4 selected 40730-like and 5 selected RlmN-like protein sequences. Amino acid coloring reflects physicochemical properties. Conserved residues within or across two protein families are boxed. Dashes indicate gaps in the sequence alignment. Protein IDs used for alignment were listed in Table S8.

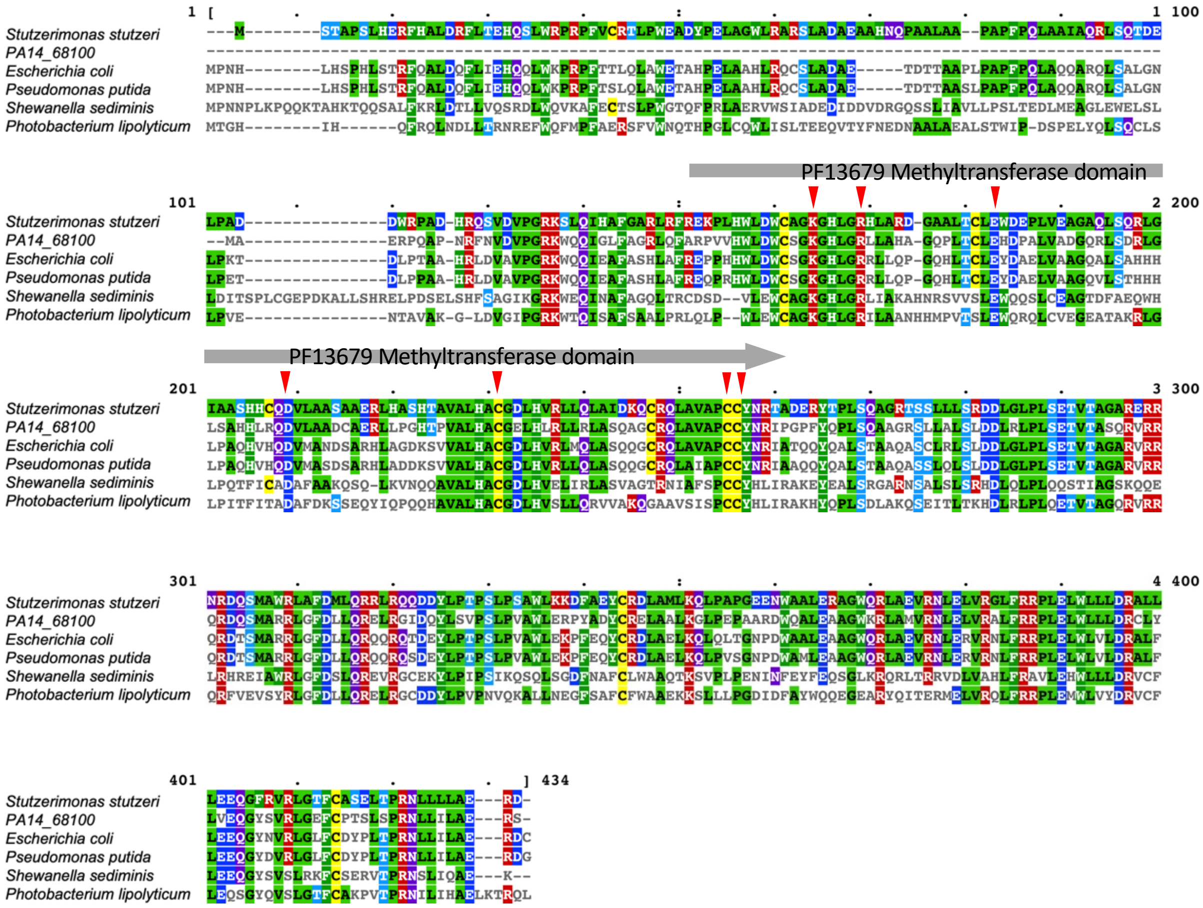

**Supplementary Figure 17. Sequence alignment of PA14_68100 homologs.** Sequences were retrieved from Uniprot using BLASTp. Amino acid coloring reflects physicochemical properties. Conserved residues in the proximity of the RNA and SAM substrate are indicated by triangles. Protein IDs used for alignment were listed in Table S8.

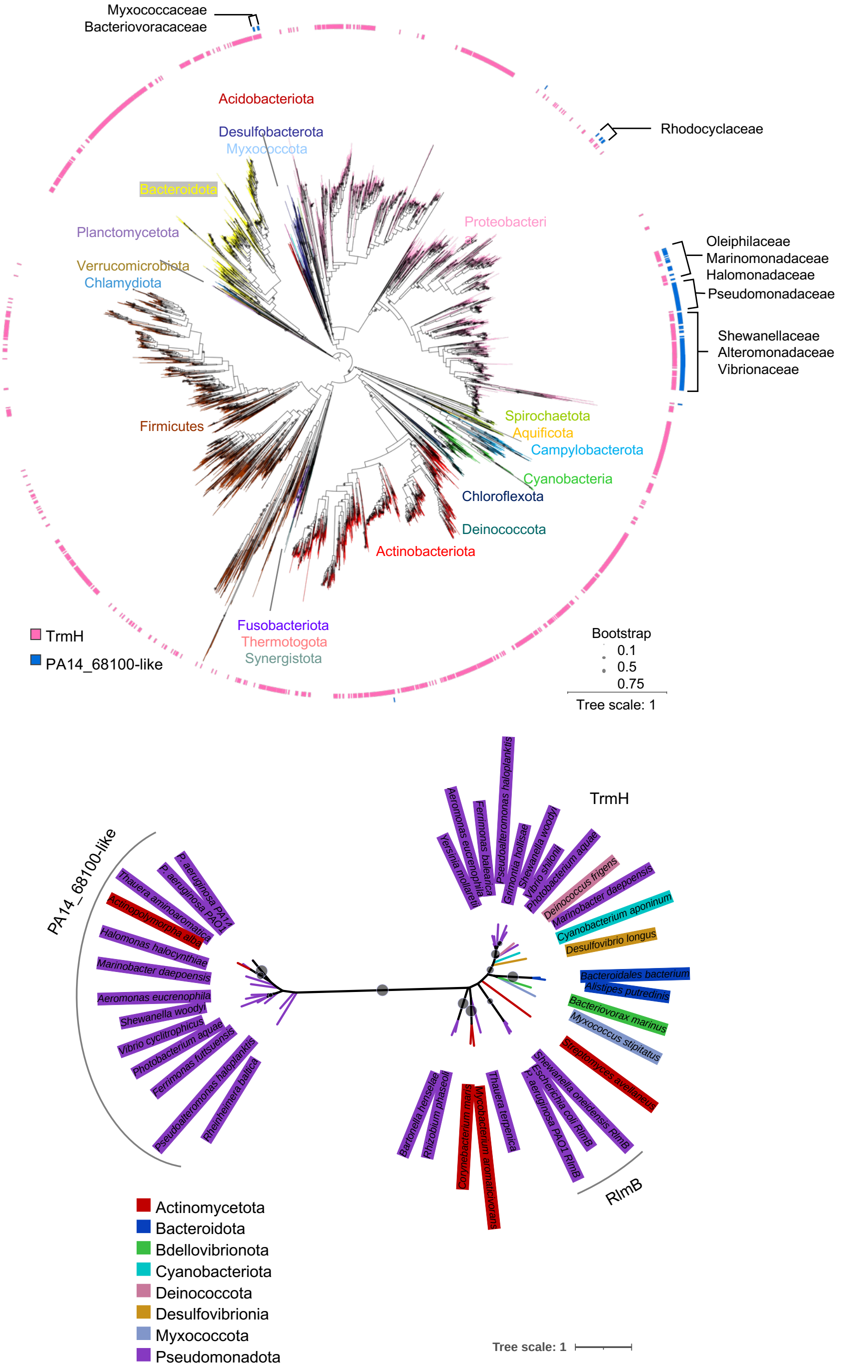


**Supplementary Figure 18. Taxonomic and phylogenetic analysis of TrmH and PA14_68100-like proteins. (A)** Distribution of TrmH and PA14_68100-like proteins in representative bacterial genomes. A maximum likelihood tree of 10 concatenated ribosomal proteins was created for the 6,616 complete representative genomes in the BV-BRC database (https://www.bv-brc.org/) as collected in Jan 2021. The branches are colored by phyla. The presence of TrmH (red) and PA14_68100-like proteins (blue) were noted in the outside circles. The families that encode PA14_68100-like proteins are indicated in the outer circle. The branches with bootstrap support value less than 0.75 were indicated by dots. For better visualization, phyla with few leaves are not annotated. (**B**) A maximum-likelihood phylogenetic tree of TrmH, RlmB and PA14_68100 like proteins. Sequences are obtained by BLASTp search in 6616 representative bacterial genomes. Branches are colored by phylogenetic affiliation at phylum level. Branches with less 0.5 bootsrap value are indicated by dots.

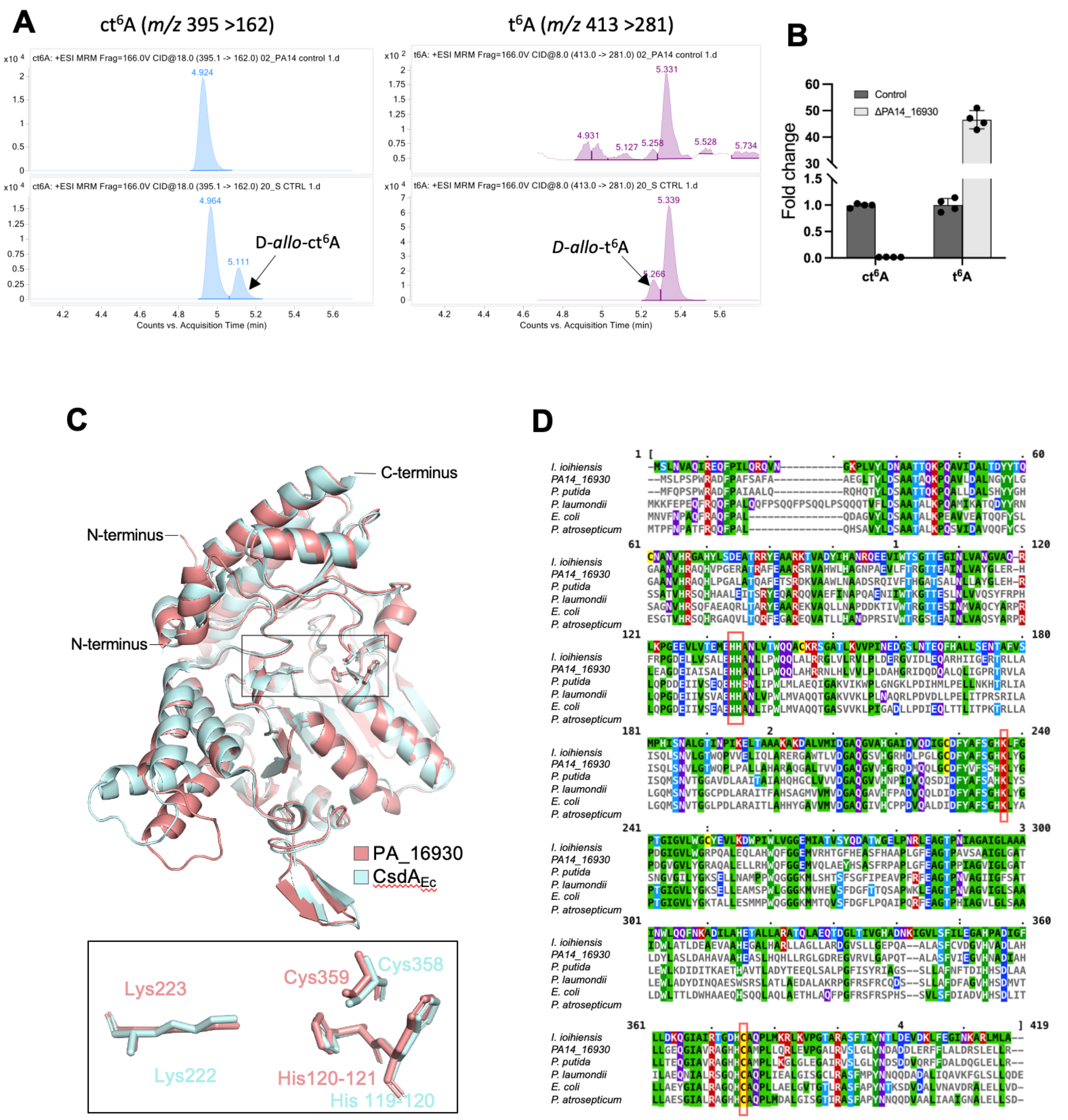


**Supplementary Figure 19. Annotation of PA14_16930 as CsdA involved in ct$^6$A formation**. (**A**) Identification of ct$^6$A and t$^6$A is complicated by RNA processing artifacts. Extracted ion chromatograms of ct$^6$A and t$^6$A in tRNA hydrolyzed in basic Tris-containing buffer (lower panel) and neutral Tris-free conditions. The isolation and hydrolysis of tRNA were performed under acidic and neutral Tris-free conditions following the protocol of Miyauchi *et al.*{Miyauchi, 2013 #5976}. (**B**). Analysis of ct$^6$A and t$^6$A levels in total RNA from the CsdA (PA14_16930) mutant strain using neutral RNA hydrolysis protocol. The fold-change values were calculated relative a transposon intergenic mutant control (n=4). (**C**) Alignment of predicted structure of PA14_16930 and crystal structure of CsdA in *E. coli* (PDB 5FT4). Three conserved residues in the sulfur transition are depicted in the box. (**D**) Sequence alignment of PA14_16930 and CsdA proteins. The conserved residues in sulfur transition are boxed. Uniport ID: *I. ioihiensis*, Idiomarina loihiensis, Q5QXG2, *P. putida*, *Pseudomonas putida*, Q9Z408, *P. laumondii*, *Photorhabdus laumondii*, Q7N8R7, *E. coli*, Q46925, *P. atrosepticum*, *Pectobacterium atrosepticum*, Q6D8G2.

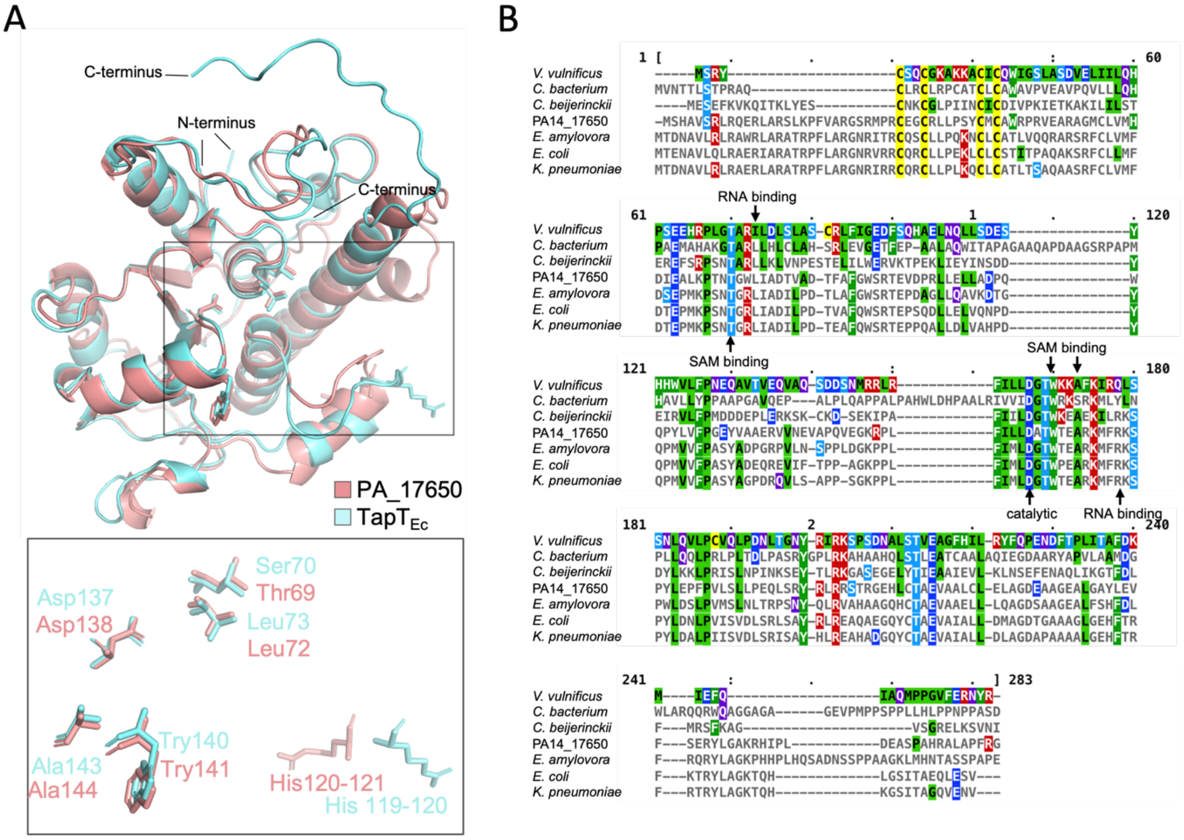


**Supplementary Figure 20. The alignment of PA14_17650 and TapT$_{Ec}$.** (A) The structural alignment of predicted structure of PA14_17650 and predicted structure of TapT in *E. coli* (Uniport Q47319). The stick cartoon represent the conserved residues in the sulfur transition (boxed on right). (B) The sequence alignment of PA14_17650 and TapT proteins. The conserved residues in substrate binding are indicated. UniprotID: *E. coli*, Q47319, *Erwinia amylovora*, D4HZ44, *Vibrio vulnificus*, Q7MMG0, *Klebsiella pneumoniae*, W1DMM7, *Clostridium beijerinckii*, Q549D3, *Comamonadaceae bacterium*, A0A4Q3M049.

### *Supplementary Tables*

**Supplementary Table 1. Configuration of Tecan EVO150 parameters used for magnetic beads-based tRNA isolation from crude lysates.** *The tips for wash buffer aliquot are re-cycled, so the dispense "move tips to left" function is not used to avoid potential contamination. #Tips for supernatant removal are also used for beads wash after tip wash, so the "move tips to left" function is not used in "Tip wash" step to avoid contamination in Waste plate. &Command "Beads wash buffer residue removal" was repeated twice, and the interval between each command is 1 min to ensure complete removal of ethanol.

| Liquid transfer steps | Aspirate parameters | | | Dispense parameters | | | Repeat |
|---|---|---|---|---|---|---|---|
| | Height (mm) | Speed (μL/s) | System trailing airgap (μL) | Height (mm) | Speed (μL/s) | Move tips to left (Y/N) | |
| Cell lysate transfer | 0.1 | 8 | 2 | -0.5 | 15 | Y | - |
| Cell lysate and Binding buffer-1 mix | -0.5 | 50 | - | -1.5 | 20 | Y | 5 |
| Wash buffer aliquot* | -0.5 | 45 | - | -1.0 | 110 | N | 10 |
| 1st supernatant transfer | 0 | 8 | 2 | -0.5 | 15 | Y | - |
| 1st residue removal | 0.1 | 8 | 3 | 1.5 | 15 | Y | 2 |
| Tip wash# | -0.5 | 50 | - | -1.5 | 200 | N | - |
| Beads wash | -0.5 | 45 | - | -1.0 | 110 | Y | 2 |
| Wash buffer residue removal& | 0.6 | 8 | 2 | -0.5 | 15 | Y | 2 |
| 2nd supernatant transfer | -1.5 | 50 | - | -1.5 | 20 | Y | - |
| RNA elution | 0.1 | 8 | 2 | -0.2 | 40 | Y | - |
| Supernatant discard | 0.1 | 8 | 2 | -1.5 | 20 | Y | 3 |

**Supplementary Table 3.** Detect of limit (LOD) and Quantification of limit (LOQ) of modified ribonucleosides using the rapid UHPLC/MS method developed in this study. These modified ribonucleosides were detected in the presence of 1000-fold excess of canonical ribonucleosides (rA, rC, rU and rG) to account for the signal suppression effect during real sample analysis.

| Modifications | Intercept | Std. Error (σ) | Slope (S) | $R^2$ | LOD (fmol) | LOQ (fmol) |
|---|---|---|---|---|---|---|
| **f5C** | 87.54 | 193.2 | 2999 | 0.9996 | 0.19 | 0.64 |
| **hm5C** | 72.89 | 67.86 | 658.9 | 0.9994 | 0.31 | 1.03 |
| **I** | 131.7 | 190.5 | 9599 | 0.9992 | 0.06 | 0.2 |
| **m5C** | 64.33 | 137.5 | 916.4 | 0.9978 | 0.45 | 1.5 |
| **m6A** | 217 | 76.39 | 2165 | 0.9974 | 0.11 | 0.35 |
| **m6Am** | -28.93 | 83.92 | 3564 | 0.9989 | 0.07 | 0.24 |
| **mcm5s2U** | -816.8 | 331.5 | 478.4 | 0.9995 | 2.08 | 6.93 |
